# Spatial genomics of AAVs reveals mechanism of transcriptional crosstalk that enables targeted delivery of large genetic cargo

**DOI:** 10.1101/2023.12.23.573214

**Authors:** Gerard M. Coughlin, Máté Borsos, Nathan Appling, Bre’Anna H. Barcelona, Acacia M. H. Mayfield, Elisha D. Mackey, Rana A. Eser, Xinhong Chen, Sripriya Ravindra Kumar, Viviana Gradinaru

## Abstract

Integrating cell type-specific regulatory elements (e.g. enhancers) with recombinant adeno-associated viruses (AAVs) can provide broad and efficient genetic access to specific cell types. However, the packaging capacity of AAVs restricts the size of both the enhancers and the cargo that can be delivered. Transcriptional crosstalk offers a novel paradigm for cell type-specific expression of large cargo, by separating distally-acting regulatory elements into a second AAV genome. Here, we identify and profile transcriptional crosstalk in AAV genomes carrying 11 different enhancers active in mouse brain. To understand transcriptional crosstalk, we develop spatial genomics methods to identify and localize AAV genomes and their concatemeric forms in cultured cells and in tissue. Using these methods, we construct detailed views of the dynamics of AAV transduction and demonstrate that transcriptional crosstalk is dependent upon concatemer formation. Finally, we leverage transcriptional crosstalk to drive expression of a large Cas9 cargo in a cell type-specific manner with systemically-administered engineered AAVs and demonstrate AAV-delivered, minimally-invasive, cell type-specific gene editing in wildtype animals that recapitulates known disease phenotypes.

**Highlights:**

- Transcriptional crosstalk between enhancers and promoters delivered in *trans* by AAVs is a generalized phenomenon.
- Spatial genomics techniques, AAV-Zombie and SpECTr, reveal that AAV genome concatemerization facilitates transcriptional crosstalk.
- Transcriptional crosstalk can be leveraged for minimally-invasive, targeted AAV delivery of large cargo, including machinery for CRISPR-based gene editing and manipulation.
- Transcriptional crosstalk enables cell-type specific gene disruption in wildtype animals, recapitulating behavioural phenotypes of genetic knockouts.

## Introduction

Recombinant adeno-associated viruses (AAVs) are versatile tools for transfer of genetic material, capable of transducing both dividing and non-dividing cells, with minimal immunogenicity^1–5^. Maintenance of the AAV genome as circular monomeric or concatemeric episomes provides long-term expression^6–10^. The tropism of AAVs can be altered by modifying residues on the AAV capsid surface and directed evolution has yielded a toolkit of capsids with diverse tropisms, including variants that can efficiently and broadly transduce target organs following systemic administration^11–23^.

AAV transduction can also be directed through inclusion of regulatory elements, including enhancer sequences mined from the host genome. Advances in single-cell epigenomics and transcriptomics have facilitated identification of cell type-specific enhancers that, when transplanted into AAV genomes, can drive expression in a cell type-specific manner^24–33^. Pairing such regulatory elements with engineered capsids offers the potential for genetic access to specific cell types without the need for transgenic driver lines, opening avenues for targeted manipulation in unconventional model organisms and in translational contexts. Integrating enhancer-driven expression with genome editing and manipulation using CRISPR (clustered regularly interspaced short palindromic repeats)-Cas (CRISPR associated)-based tools^34–36^ can facilitate understanding of gene function in targeted cell types without confounds due to on- or off-target editing in other cell types.

AAV delivery of enhancer-driven CRISPR-Cas systems is hindered by AAVs’ relatively low packaging capacity of 4.7 kb, including the requisite inverted terminal repeats (ITRs), and large size of enhancers and CRISPR effector proteins. In the host genome, enhancers and their target gene(s) are often separated by large distances; chromatin looping can bring distal enhancers and promoters closer in proximity^37,38^. Similarly, Duan and colleagues^39^ demonstrated that a ubiquitous enhancer in one AAV genome can increase expression from another AAV genome when delivered in *trans*. This phenomenon (which we term transcriptional crosstalk) represents an underexplored approach for delivery of large cargo to specific cell types.

Increased understanding of AAV genome processing and genome-genome interaction at a single cell level would facilitate implementation of transcriptional crosstalk as a large cargo delivery method. Although existing techniques to visualize AAV genomes *in situ*^40,41^ can provide valuable subcellular information about AAV transduction, such methods are not able to detect certain endpoints of AAV genome processing (e.g. concatemeric episomes). Askary and colleagues^42^ recently developed the Zombie method, in which phage polymerase promoters and barcodes are incorporated into the DNA of interest. Phage RNA polymerase added to the fixed tissue transcribes the barcode, yielding RNA transcripts that can be detected through high sensitivity HCR-FISH (hybridization chain reaction fluorescence *in situ* hybridization)^43^. These transcripts serve as proxies for the encoding genome. Importantly, the use of enzymatic amplification may enable specific detection of certain AAV genome states *in situ*.

Here we investigate transcriptional crosstalk, demonstrating its generalizability to a broad array of cell type-specific enhancers. Using novel single-molecule spatial genomics methods based on Zombie, we explore the mechanism of transcriptional crosstalk *in vitro* and *in vivo* and demonstrate critical roles for AAV concatemers in facilitating this phenomenon. Finally, we leverage transcriptional crosstalk to achieve cell type-specific delivery of a large Cas9 cargo, following systemic injection of an engineered AAV, resulting in targeted genome editing that recapitulates known behavioural phenotypes.

## Results

### Broad transcriptional crosstalk between enhancers and promoters delivered in separate AAV genomes

Transcriptional crosstalk between AAV genomes can occur when regulatory elements in one genome interact with those of another. The Ple155 element^44^ drives strong expression in mouse cerebellar Purkinje cells (PCs) following systemic delivery via a blood-brain barrier (BBB)-penetrant engineered AAV (AAV-PHP.eB^12^). Conversely, the mDLX enhancer^24^ paired with a minimal beta-globin promoter (mDLX-minBG) directs expression to forebrain interneurons, but not PCs. However, following co-transduction of these viruses, we observed strong expression of the mDLX-minBG-driven transgene in PCs (Fig. 1a,b and Extended Data Fig. 1a). This result suggests that elements in the Ple155 sequence can interact with elements in the mDLX-minBG genome and increase expression of the latter in a cell type-specific manner.

**Figure 1.**
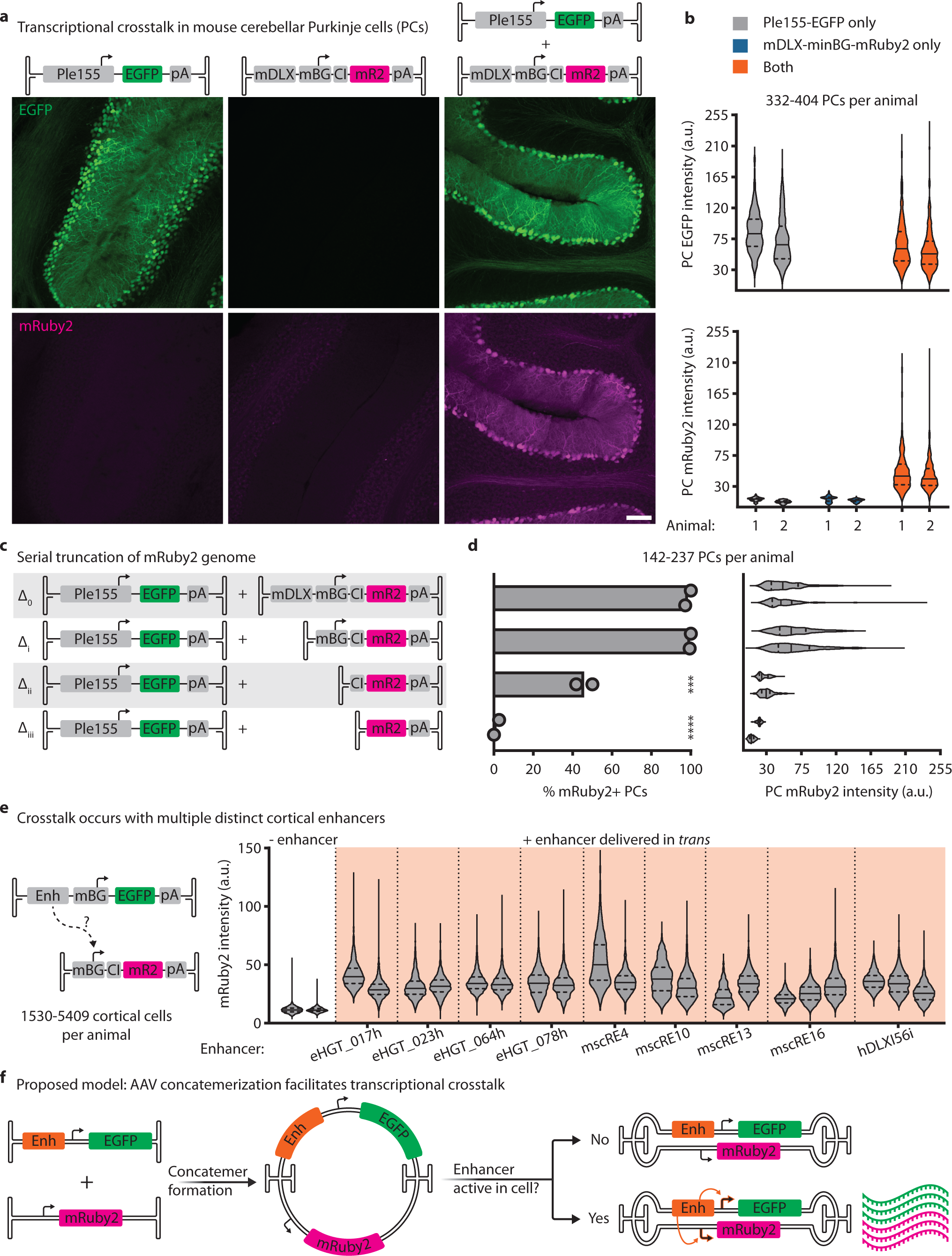
Broad transcriptional crosstalk between enhancers and promoters delivered in separate AAV genomes. **a**, Transcriptional crosstalk. Left column: when injected alone, the AAV-delivered Ple155 element directs strong expression to cerebellar Purkinje cells (PCs). Middle: AAV-delivered mDLX-minBG-driven mRuby2 does not yield any detectable PC transduction. Right: co-administration of both AAVs results in unexpected mRuby2 expression in PCs. All genomes delivered at 1e12 vg dose in AAV-PHP.eB. Scale bar = 100 μm. **b**, Distribution of PC cell body EGFP (top) and mRuby2 (bottom) intensities from animals shown in (**a)**. Each violin plot represents EGFP or mRuby2 intensity from PCs in one animal. *n* = 2 animals per condition. **c**, Schematic of serially-truncated mDLX-minBG-mRuby2 constructs, coinjected with Ple155-EGFP, to assess necessity of elements for transcriptional crosstalk. **d**, Quantification of results for truncation conditions shown in (**c**), quantified as percent of PCs positive for mRuby2 (left) and PC mRuby2 fluorescence intensity (right). Bars represent mean, with statistical significance determined by one-way analysis of variance and Dunnett’s multiple comparison test against full-length mRuby2 genome. Each violin plot represents mRuby2 intensity from PCs in one animal. *n* = 2 animals per condition. All genomes delivered at 5e11 vg dose in AAV-PHP.eB. **e**, Screen of 9 cortical enhancers for ability to upregulate expression of minBG promoter-driven mRuby2 delivered in *trans*. Each violin plot represents mRuby2 intensity from mRuby2+ cortical cells in one animal. *n* = 2 animals per condition, except mscRE16 and hDLXI56i, in which *n* = 3. All genomes delivered at 1e12 vg dose in AAV-PHP.eB. **f**, Proposed model for transcriptional crosstalk. Formation of concatemeric episomes places enhancer and promoter elements that were delivered in *trans* into a *cis* conformation. This concatemerization facilitates interaction of the enhancer with the promoter that was delivered in *trans*, resulting in increased expression in cells where the enhancer is active. For violin plots, the solid line is the median, and upper and lower dashed lines are quartiles. CI = chimeric intron. \*\*\**P* < .001, \*\*\*\**P* < .0001.

To identify which elements in the mDLX-minBG sequence are necessary for this crosstalk, we serially truncated the mDLX-minBG genome (Fig. 1c,d and Extended Data Fig. 1b,c). Removal of the mDLX enhancer did not produce a detectable effect on crosstalk (truncation Δ_i_), whereas removal of the minBG promoter decreased both the percent of mRuby2-positive PCs and the PC mRuby2 intensity (truncations Δ_ii_ and Δ_iii_). These data point to a model in which elements in the Ple155 interact with the minBG promoter, reminiscent of the classical description of enhancer-promoter interaction^37,38^.

Given this model for transcriptional crosstalk, we expect to observe this behaviour with multiple enhancer sequences. Thus, we screened a panel of 9 characterized cortical enhancer sequences^24,26,27^, using a minBG-driven mRuby2 crosstalk reporter virus (Fig. 1e and Extended Data Fig. 2a-c). In all 9 cases, presence of the enhancer resulted in an increase in expression from the reporter genome delivered in *trans*, when compared to a ‘no enhancer’ condition (Fig. 1e).

To further demonstrate the generalized nature of transcriptional crosstalk, we used the ubiquitous cytomegalovirus immediate-early enhancer^45^ (CMVe) and the super core promoter 1 (SCP1)^46^ in combination with a cocktail of BBB-penetrant and peripheral nervous system-tropic engineered AAV capsids (AAV-PHP.eB and AAV-MaCPNS2^15^), to provide broad central and peripheral nervous system coverage (Extended Data Fig. 2d). We observed increased tdTomato crosstalk reporter expression in cerebellum, proximal colon, dorsal root ganglia, and liver with an enhancer delivered in *trans* vs. the no enhancer condition.

These results support a generalized model for transcriptional crosstalk, in which enhancer elements on one AAV genome can interact with and drive expression from a promoter on another AAV genome. As this interaction is more likely to occur between elements in *cis,* we and others^39^ propose that concatemerization of AAVs could enable transcriptional crosstalk, by placing elements delivered in *trans* into a *cis* conformation (Fig. 1f).

### AAV-Zombie reveals intracellular AAV genome localization in cultured cells and in tissue

To further explore transcriptional crosstalk, we required methods for single molecule AAV genome localization in intact cells and tissue. We therefore adapted the Zombie method^42^, by incorporating phage RNA polymerase promoters and barcodes into the AAV genome (Fig. 2a). *In situ* transcription and HCR-FISH against the nascent barcoded transcript allow for subcellular localization of both single-stranded AAV (ssAAV) and self-complementary AAV (scAAV) genomes (Fig. 2b).

**Figure 2.**
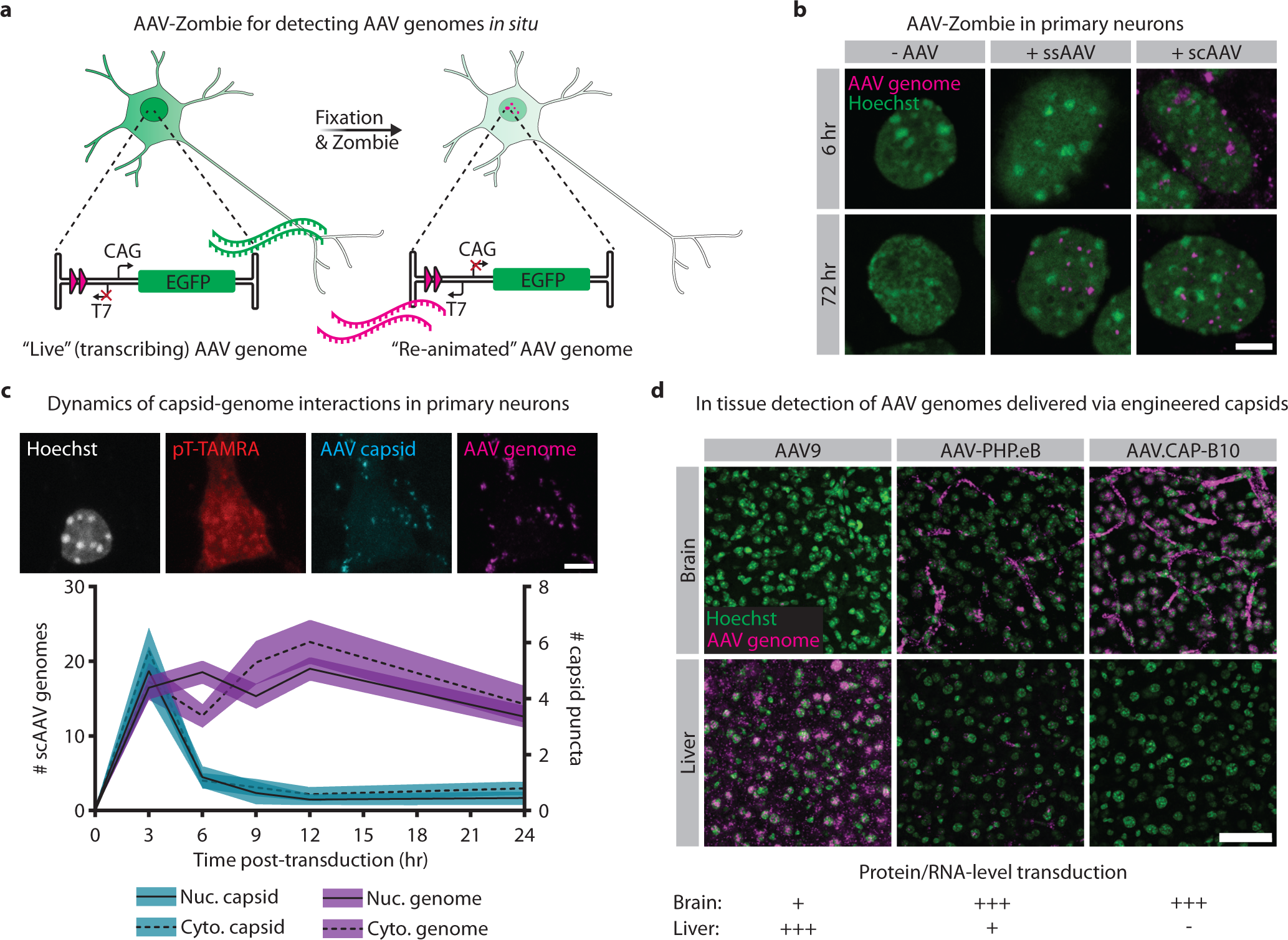
AAV-Zombie reveals intracellular AAV genome localization in cultured cells and in tissue. **a**, Schematic of AAV-Zombie. A barcode and phage RNA polymerase promoter are integrated into the AAV genome. While the cell is alive, the barcode is not transcribed. After fixation, *in situ* transcription of the barcode by phage RNA polymerase yields barcoded transcripts that can be detected by HCR-FISH. These transcripts serve as a proxy for the AAV genome. **b**, Detection of single-stranded and self-complementary AAV genomes (ssAAV and scAAV, respectively) in cultured primary neurons. At 6 hours post-transduction, ssAAV genomes are rarely detected due to the necessity of second strand synthesis, whereas scAAV genomes are readily detected in and outside the nucleus. At 72 hours, genomes of both formats are detected in the nucleus. All genomes delivered at 1e5 MOI in AAV6. Scale bar = 5 μm. **c**, Time course of AAV capsids and scAAV genomes in nucleus and cytoplasm of primary neurons. Capsids were detected through immunofluorescence with an antibody against linear epitopes. Cytoplasm was labeled with a TAMRA-conjugated polyT probe. Genomes were delivered at 1e6 MOI in AAV-DJ. Black line is mean; shaded area is 95% confidence interval. *n* = 243 (t = 0 hr), 191 (3 hr), 317 (6 hr), 212 (9 hr), 220 (12 hr), 255 (24 hr) neurons per time point. Scale bar = 5 μm. **d**, AAV-Zombie detection of AAV genomes in mouse brain and liver 1 day post-injection, following systemic delivery by AAV9, AAV-PHP.eB, or AAV.CAP-B10, at 3e11 vg dose. Distribution of AAV genomes recapitulates known protein- and RNA-level transduction profiles (bottom). Representative images from *n* = 3 animals per condition. Scale bar = 50 μm.

Understanding AAV trafficking and processing at early stages of transduction can provide invaluable insights into the vector’s biology. To investigate the dynamics of AAV capsid-genome interaction, we paired AAV-Zombie with immunohistochemistry (IHC) and profiled transduction in primary neuron culture over 24 hours (Fig. 2c and Extended Data Fig. 3a-c). As expected, capsid puncta were transient, in both the cytoplasm and nucleus, peaking early in transduction and dropping back to baseline by 12 hours. scAAV genomes were more stable over time in both compartments. Importantly, more than 96% of capsid puncta colocalized with a genome (across all time points); the fraction of genome puncta colocalizing with a capsid was lower and decreased over time (Extended Data Fig. 3b,c).

Given these promising results of AAV-Zombie in cultured cells, we then tested its performance in mouse brain and liver, comparing the DNA-level transduction between two generations of engineered capsids and their parent AAV9 (Fig. 2d and Extended Data Fig. 3d). Consistent with known protein- and RNA-level transduction patterns^12,14,18^ (Fig. 2d, bottom), AAV9 strongly transduced the liver, but was rarely observed in the brain, while AAV-PHP.eB and AAV.CAP-B10^14^ both transduced the brain well, with reduced liver transduction for AAV-PHP.eB and no detected liver transduction for AAV.CAP-B10. These results demonstrate the power of AAV-Zombie for exploring AAV transduction, both in cultured cells and in tissue.

### SpECTr reveals spatiotemporal dynamics of AAV concatemerization

To enable detection of concatemerized AAV genomes, we adapted AAV-Zombie by separating the phage polymerase promoter and barcode (ConcBC) into separate AAV genomes (termed the “T7/SP6” and “barcode” genomes, respectively) (Fig. 3a). Concatemerization of these two genomes orients the T7 promoter and ConcBC such that T7 polymerase can transcribe the barcode. The T7/SP6 genome also contains a barcode (GenBC) driven by an SP6 RNA polymerase promoter, allowing detection of that AAV genome independent of concatemerization. The short length of the phage promoters and barcodes (∼20 nt and 100-250 nt, respectively) leaves ample space for strong mammalian promoters and reporter genes. Thus, following cotransduction, fixation, and Zombie, we could detect the concatemer-independent barcode, concatemer-dependent barcode, as well as reporter gene transcripts (Fig. 3b), providing single-molecule information about AAV transduction, concatemer formation, and expression in single cells. We term this method SpECTr, for “<u>Sp</u>ECTr <u>E</u>nables AAV <u>C</u>oncatemer <u>Tr</u>acking”.

**Figure 3.**
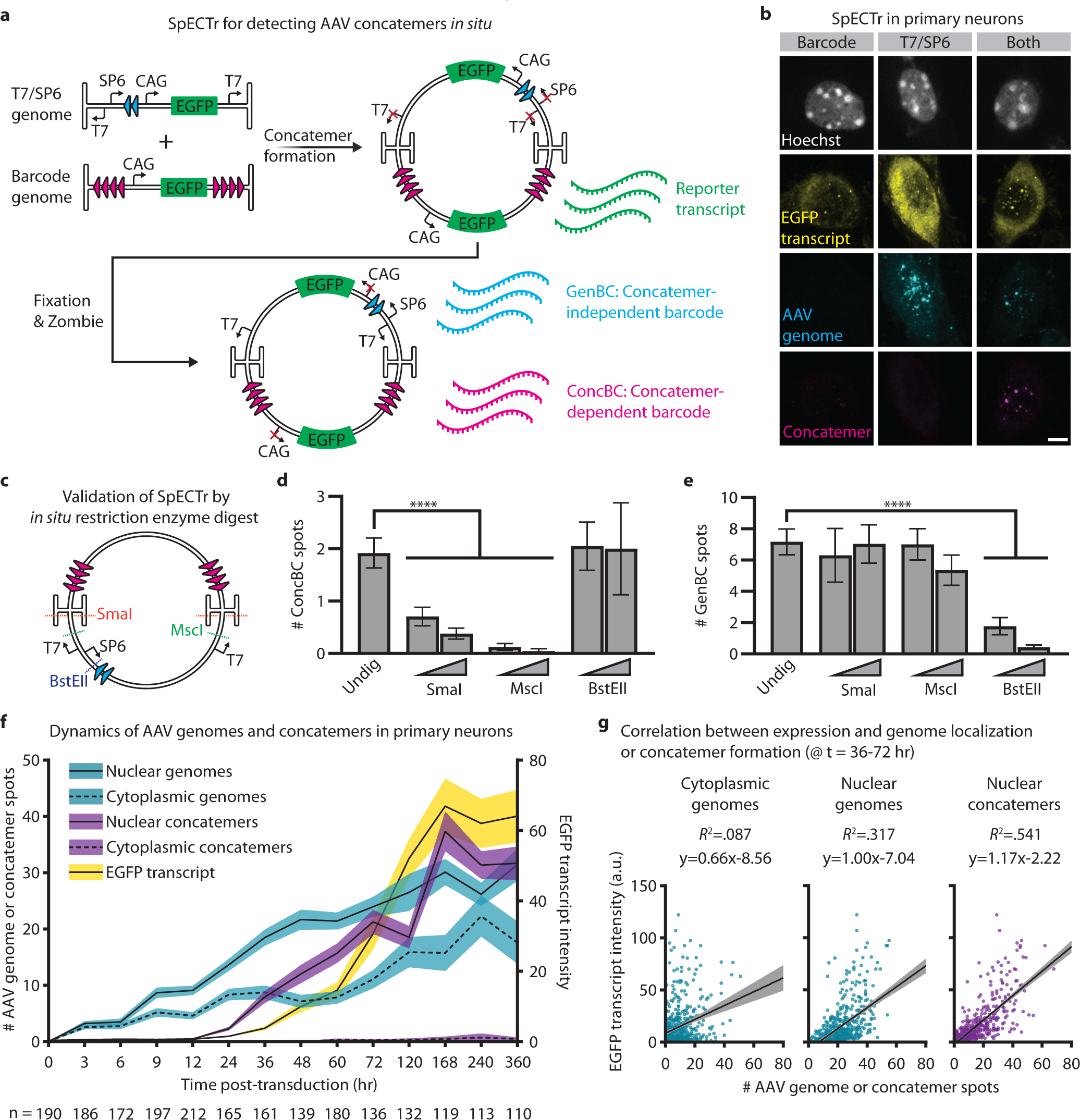
SpECTr reveals spatiotemporal dynamics of AAV concatemerization. **a**, Schematic of SpECTr. Two AAV genomes are used: the T7/SP6 genome delivers the phage RNA polymerase promoters, and the barcode genome delivers a concatemerization-dependent barcode (ConcBC). Concatemerization of these two genomes orients the T7 promoter and the ConcBC such that T7 RNA polymerase can transcribe the ConcBC. The T7/SP6 genome also contains a concatemerization-independent barcode (GenBC), driven by an SP6 RNA promoter in *cis*. Both genomes carry a CAG-driven EGFP, to enable readout of transgene expression. **b**, Specificity of SpECTr in primary neurons collected 72 hours post-transduction. ConcBC spots are only detected in cells co-transduced by both genomes (right column). Scale bar = 5 μm. **c-e**, Validation of SpECTr through *in situ* restriction enzyme digest of HEK293T cells transduced with SpECTr genomes. **c**, Map of model AAV concatemer containing 1 copy of the T7/SP6 genome and 1 copy of the barcode genome, showing SmaI, MscI and BstEII restriction enzyme sites. **d,e**, Number of ConcBC spots (**d**) and GenBC spots (**e**) detected following *in situ* restriction enzyme digests, with low (20 U/mL) and high (200 U/mL) restriction enzyme concentrations. “Undig”: undigested condition in which fixed cells were incubated at 37 °C in restriction enzyme buffer, without any restriction enzyme present. Statistical significance was determined using Kruskal-Wallis test against the undigested condition. Bars are mean ± s.e.m. *n* = 138 (Undigested), 99 (low SmaI), 89 (high SmaI), 87 (low MscI), 22 (high MscI), 40 (low BstEII), 26 (high BstEII) cells per condition. **f**, Time course of AAV transduction, concatemer formation, and EGFP reporter transcription in primary neurons. Cytoplasm was labeled with a TAMRA-conjugated polyT probe, and nucleus with Hoechst. EGFP transcript intensity was quantified in entire cell body; AAV genomes and concatemers were quantified in nucleus and cytoplasm separately. Black line is mean; shaded area is 95% confidence interval. Number of neurons per time point is indicated on figure. **g**, Correlation between EGFP reporter expression and cytoplasmic genome spots, nuclear genome spots, and nuclear concatemer spots. *n* = 616 primary neurons, pooled from t = 36, 48, 60, and 72 hr time points. For all experiments, genomes were delivered at 1e6 MOI in AAV-DJ. \*\*\*\**P* < .0001.

To confirm that the ConcBC transcript arises from a single molecule containing both the T7 promoter and ConcBC, we performed *in situ* restriction enzyme digests on AAV-DJ-transduced and fixed HEK293T cells before barcode transcription. Digestion with SmaI (which cuts within the AAV ITR) or MscI (which cuts immediately downstream of the T7 promoter) significantly reduced the number of detected ConcBC spots, without affecting the number of GenBC spots. Conversely, digestion with BstEII (which cuts immediately downstream of the SP6 promoter) significantly reduced the number of GenBC spots without affecting the number of ConcBC spots (Fig. 3c-e and Extended Data Fig. 4). These results provided confidence that SpECTr specifically detects AAV concatemers *in situ*.

To test the utility of SpECTr for exploring AAV transduction, we conducted a time course of AAV-DJ transduction in primary neurons, collecting samples at 14 time points over 360 hours post-transduction (Fig. 3f and Extended Data Fig. 5a). As expected, we observed an immediate and steadily increasing count of AAV genomes in both the nucleus and cytoplasm. Nuclear concatemeric genome counts began to rise between 12 and 24 hours post-transduction, followed shortly after by EGFP transcript intensity. The relative order of these increases (genomes, concatemers, transcript) further supports that SpECTr is detecting AAV concatemers, and that AAV concatemerization is important for strong transgene expression. Consistent with specific detection of AAV concatemers, cytoplasmic concatemer counts were low at all time points measured (mean < 1 and median = 0, per cell, for each time point).

SpECTr provides subcellular and multiparametric data about AAV transduction, enabling us to explore relationships between genome forms, their localization, and expression at the single-cell level (Fig. 3g and Extended Data Fig. 5b,c). Notably, we observed a weak correlation of reporter transcript intensity with cytoplasmic genome counts (*R*^2^ = .087), a moderate correlation with nuclear genome counts (*R*^2^ = .317), and a strong correlation with nuclear concatemer counts (*R*^2^ = .541) (Fig. 3g).

These results establish AAV-Zombie and SpECTr as validated tools for visualizing AAV genomes and concatemers *in situ*.

### Reducing AAV concatemer formation decreases transcriptional crosstalk between AAV genomes

Using SpECTr to visualize AAV concatemers, we next explored the mechanistic connection between concatemerization and transcriptional crosstalk. If concatemerization of AAV genomes enables transcriptional crosstalk (Fig. 1f), then we expect reductions in concatemer formation to reduce transcriptional crosstalk. We first tested this hypothesis in HEK293T cells with the ubiquitous CMVe, comparing AAV-DJ transduction to plasmid transfection (Extended Data Fig. 6). As expected, transcriptional crosstalk was apparent following cotransduction by AAVs, but not after cotransfection of the corresponding genome plasmids. Transfection of a “plasmid concatemer”, consisting of the entire tdTomato-containing genome inserted outside the ITRs of the TagBFP-containing genome plasmid, recapitulated the co-transduction result. These data suggest that transcriptional crosstalk occurs when the enhancer and promoter are in a *cis* conformation.

We next tested this hypothesis *in vivo.* Previous research has implicated DNA repair pathways in recognizing and processing free ITR ends, resulting in formation of concatemeric AAV episomes^47–53^. In particular, *Prkdc^scid/scid^*mice (hereafter referred to as SCID mice), which have a loss of function in the DNA double-strand break repair enzyme Prkdc, show reduced concatemer formation in bulk muscle^48^ and liver^50^, and lower expression from concatemerization-dependent AAVs^47^. However, neither concatemer formation at a single-cell level nor transcriptional crosstalk have been explored in SCID mice.

We first validated SpECTr for detection of AAV concatemers in tissues, including cortex, liver, and cerebellum (Extended Data Fig. 7a). Then, to enable paired measurement of concatemer formation and transcriptional crosstalk in the same animals, we integrated SpECTr components into the Ple155 and mDLX-minBG AAV genome pair. Reasoning that the high doses of AAVs we used previously (1e12 vg, Fig. 1a) may yield many large indistinguishable spots and thus confound accurate measurement of concatemers, we injected 3e11 total vg into C57BL/6J-background SCID mice and wildtype C57BL/6J controls. Even at this reduced dose, transcriptional crosstalk was readily apparent in PCs of wildtype animals; transduction with both genomes resulted in significantly more mRuby2-positive PCs and a significant increase in PC mRuby2 intensity, compared to either single transduction condition (Fig. 4a,b and Extended Data Fig. 7b). These effects were not observed in SCID mice.

**Figure 4.**
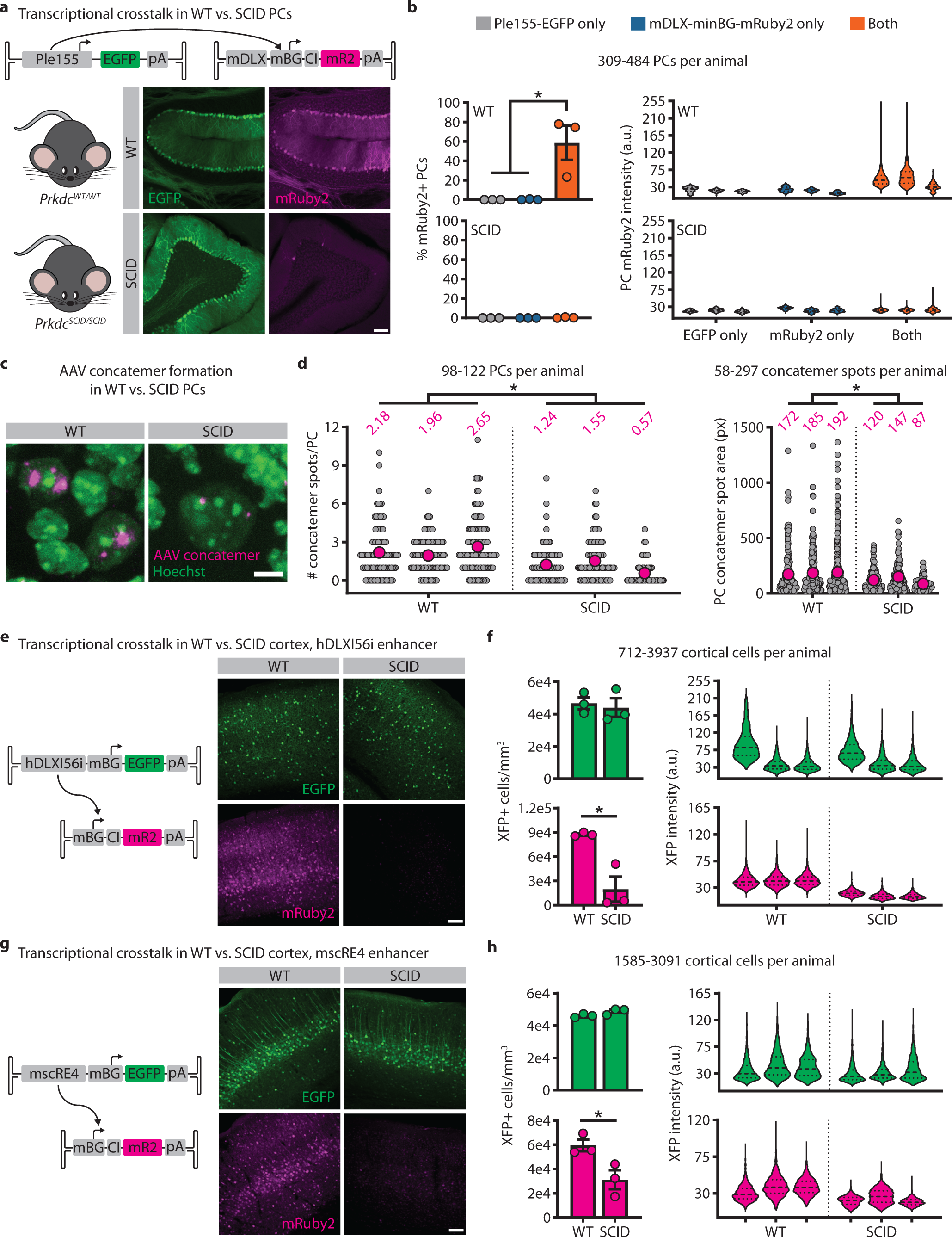
Reducing AAV concatemer formation decreases transcriptional crosstalk between AAV genomes. **a**, Representative images of transcriptional crosstalk between Ple155 and minBG promoter, in dual-injected WT and SCID mouse Purkinje cells (PCs). Both genomes delivered at 3e11 vg dose in AAV-PHP.eB. Scale bar = 100 μm. **b**, Quantification of transcriptional crosstalk shown in (**a**), comparing single injection conditions to dual injection condition, and measured as percent of PCs positive for mRuby2 (left) and PC mRuby2 fluorescence intensity (right). Statistical significance was determined using one-way analysis of variation and Tukey’s multiple comparison test. *n* = 3 animals per condition. **c**, Representative images of AAV concatemers detected with SpECTr in PCs of dual-injected WT and SCID animals shown in (**a**). Scale bar = 5 μm. **d**, Quantification of PC concatemer spot count (left) and spot size (right), in WT and SCID PCs. Each grey dot corresponds to a single PC (left) or a single concatemer spot (right). Magenta dot and number indicate mean of animal. *n* = 3 animals per condition. **e-h**, Representative images and quantification of reduced transcriptional crosstalk in SCID animals with the GABAergic interneuron enhancer hDLXI56i (**e,f**) and the layer 5 pyramidal tract neuron enhancer mscRE4 (**g,h**). Fluorescent protein (XFP) signal was amplified through IHC. Quantification is presented as number of XFP-positive cells per mm^3^ and XFP fluorescence intensity. *n* = 3 animals per condition. Scale bars = 100 μm. Bars in (**b**), (**f**), and (**h**) represent mean ± s.e.m. Statistical significance in (**d**), (**f**), and (**h**) was determined using unpaired t-tests. \**P* < .05.

To determine whether SCID PCs were deficient in AAV concatemerization, we applied AAV-Zombie and SpECTr to cerebellum sections from the same animals. PCs were identified using HCR-FISH against *Iptr1*^54^. With AAV-Zombie, we measured a 2-fold higher AAV genome count in SCID than WT PCs (Extended Data Fig. 7c,d). In a separate cohort of mice, we similarly observed significantly higher DNA-level transduction of SCID brains by AAV-PHP.eB, with no significant differences in protein-level transduction (Supplemental Fig. 1). Despite higher DNA-level transduction of SCID brains, SpECTr revealed significantly fewer and smaller ConcBC spots in PCs of SCID mice than wildtype controls (Fig. 4c,d and Extended Data Fig. 7e), indicating reduced concatemer formation in the absence of functional Prkdc.

Finally, we assessed whether the SCID mutation would affect crosstalk of other enhancers as well. We chose two additional enhancers, targeting GABAergic interneurons (hDLXI56i)^24^ and layer 5 pyramidal tract excitatory neurons (mscRE4)^26^, and coinjected these with an mRuby2 crosstalk reporter (Fig. 4e-h and Extended Data Fig. 7f,g). Consistent with our observations from the Ple155 and mDLX-minBG pair, we observed reduced transcriptional crosstalk with both enhancers in SCID mice, quantified by both number of mRuby2-positive cells per mm^3^ and fluorescence intensity of mRuby2-positive cells.

We did not detect any difference in transduction between genotypes, as assessed by number and intensity of EGFP-positive cells.

Taken together, these *in vitro* and *in vivo* results strongly suggest that AAV concatemer formation enables transcriptional crosstalk. As concatemer formation appears to be a common endpoint of AAV genome processing, and given the generalizability of the phenomenon across cell type-specific enhancers, we next explored whether we could leverage transcriptional crosstalk to achieve cell type-specific expression of large cargos.

### Transcriptional crosstalk enables all-AAV cell type-specific genome editing with CRISPR-Cas9

We reasoned that transcriptional crosstalk might enable cell type-specific delivery of larger cargo, by separating bulky gene regulatory elements from minimal promoters and coding sequences in another AAV (Fig. 5a). We explored the feasibility of this approach using *Staphylococcus aureus* Cas9 (SaCas9) as a large cargo and adopting a commonly-used reporter assay based on Ai14 mice (*Rosa26^CAG-LSL-tdTomato^*)^55–58^.

**Figure 5.**
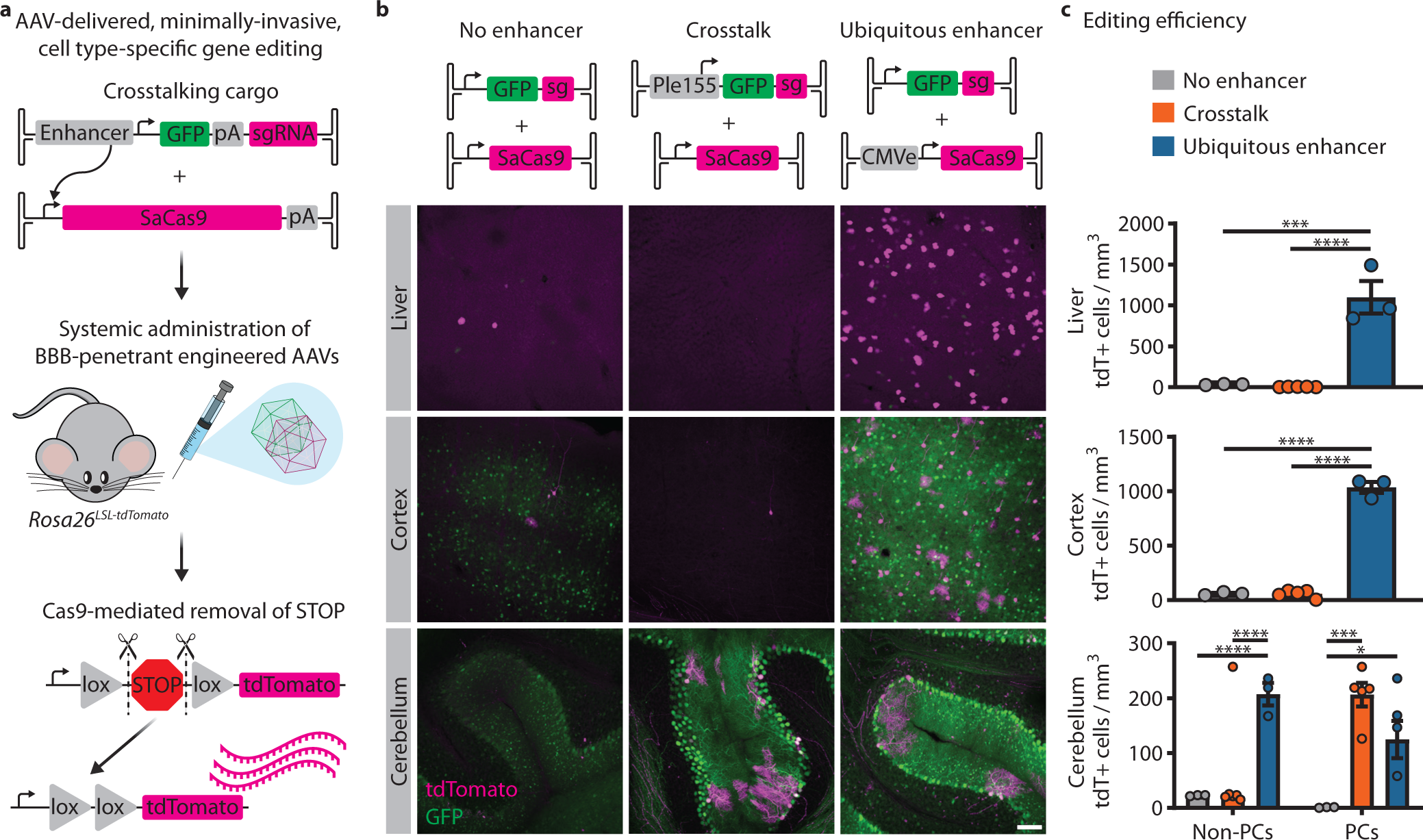
Transcriptional crosstalk enables all-AAV cell type-specific genome editing with CRISPR-Cas9. **a**, Schematic of AAV-delivered, minimally-invasive, cell type-specific gene editing. *Staphylococcus aureus* Cas9 (SaCas9), packaged with minimal elements (total size 4.2 kb), is delivered with a bulky enhancer element in *trans*, resulting in upregulation of SaCas9 expression in a cell type-specific manner. As a proof of principle, we used a common reporter assay with *Rosa26^LSL-tdTomato^* mice, in which guide RNAs direct SaCas9 to remove the stop cassette, enabling tdTomato expression. All genomes delivered at 1e12 vg dose in AAV-PHP.eB. **b**, Demonstration of crosstalk-enabled gene editing using the Ple155 element to drive SaCas9 expression in Purkinje cells (PCs). As controls, we included a no enhancer condition, as well as a condition in which SaCas9 is strongly expressed by the ubiquitous CMVe delivered in *cis*. Representative images from liver, cortex, and cerebellum. Scale bars = 100 μm. **c**, Quantification of editing efficiency, assessed by number of tdTomato-positive cells per mm^3^ of tissue. PCs and non-PCs were quantified separately. Using crosstalk to drive strong SaCas9 expression specifically in PCs restricted high efficiency editing to that cell type. Statistical significance was determined using one-way analysis of variation and Tukey’s multiple comparison test. *n* = 3 (no enhancer or ubiquitous enhancer) or 5 (crosstalk) animals per condition. Bars represent mean ± s.e.m. \**P* < .05, \*\*\**P* < .001, \*\*\*\**P* < .0001

Minimal expression of SaCas9 with no enhancer resulted in low efficiency of editing in all tissues examined (Fig. 5b,c “No enhancer”). Conversely, when SaCas9 was strongly expressed with a ubiquitous enhancer (CMVe) in *cis*, we observed a strong increase in editing efficiency in tissues of interest, compared to the no-enhancer condition. We saw a 31-fold increase in editing in the liver, a 17-fold increase in the cortex, a 9-fold increase in non-Purkinje cerebellar cells (non-PCs), and a 107-fold increase in PCs (Fig. 5b,c “Ubiquitous enhancer”). Using transcriptional crosstalk to direct SaCas9 expression specifically to PCs with the Ple155 element in the companion AAV genome, we restricted efficient editing to PCs, yielding a 177-fold increase in PC editing efficiency compared to the no-enhancer condition, with no significant increases in other tissues and cerebellar cell types (Fig. 5b,c “Crosstalk”). These results establish the utility of transcriptional crosstalk for expanding the limits of AAV-based cell type-specific genome editing and manipulation.

### Transcriptional crosstalk enables efficient gene disruption of Cacna1a in Purkinje cells, resulting in ataxic phenotypes

Efficient and specific gene editing through transcriptional crosstalk from systemically-delivered AAVs offers a means to explore gene function in a cell type-specific manner. Importantly, this strategy does not rely on transgenic lines, enabling rapid and cost-effective generation of large cohorts from easily obtained wildtype animals.

To test this approach, we targeted *Cacna1a* in wildtype C57BL/6J mice, in either a ubiquitous or PC-specific manner using transcriptional crosstalk (Fig. 6a, left side). *Cacna1a* is broadly expressed in the brain, and global knockout leads to dystonia, ataxia, cerebellar degeneration, absence seizures, and early lethality^59,60^. Targeted loss-of-function in PCs leads to ataxia^61^, whereas loss-of-function in cerebellar granular cells leads to ataxia and absence seizures^62^. Forebrain-specific deletion of *Cacna1a* causes epileptiform activity as well as learning and memory deficits^63^. Thus, understanding the function of *Cacna1a* in PCs requires methods to specifically target PCs, thereby avoiding confounds due to loss-of-function in other brain regions.

**Figure 6.**
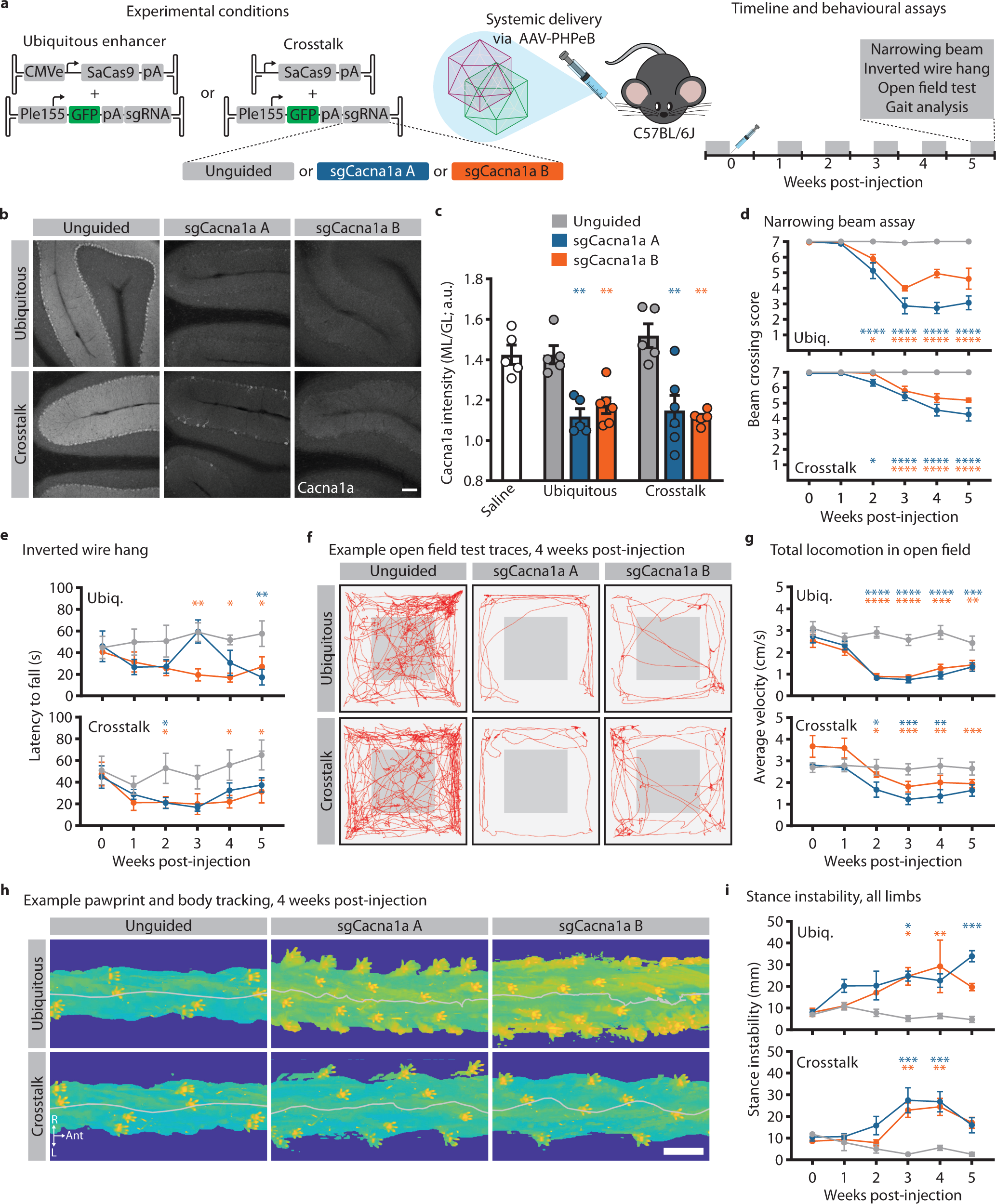
Transcriptional crosstalk enables efficient knockout of *Cacna1a* in Purkinje cells, resulting in an ataxic phenotype. **a**, Experimental design. Two conditions were tested. In the ubiquitous enhancer condition, SaCas9 was strongly expressed with CMVe delivered in *cis*. In the crosstalk condition, SaCas9 expression was restricted to Purkinje cells (PCs) through inclusion of the Ple155 element delivered in *trans.* Two sequence-independent guide RNAs targeting *Cacna1a* were used and compared to an unguided condition. All genomes delivered at 1e12 vg dose in AAV-PHP.eB. Behavioural assays were performed weekly, before and for five weeks after AAV administration. **b**, Representative images of IHC against Cacna1a in cerebellum. Scale bar = 100 μm. **c**, Quantification of Cacna1a staining in cerebellum, normalized as the intensity in the molecular layer (ML) divided by the intensity in the granular layer (GL). Statistical significance was determined by one-way analysis of variance and Dunnett’s multiple comparison tests against Cacna1a intensity in age-matched saline-injected mice. **d-i**, characterization of ataxic phenotypes following ubiquitous (“Ubiq.”, top graphs) and PC-specific (“Crosstalk”, bottom graphs) disruption of *Cacna1a*. Skilled locomotion was assessed with the narrowing beam assay (**d**), limb strength with the inverted wire hang test (**e**), locomotion in an open field (**f,g**), and gait using automated pawprint and body tracking (**h,i**). Red lines in **f** represent animal position over a 10 min trial, acquired 4 weeks post-injection. Heatmaps in **h** show pawprint positions and body tracking over a small segment of the elevated plexiglass platform used for gait analysis, at 4 weeks post-injection. Grey line indicates midline of body. Scale bar = 3 cm. Statistical significance for beam crossing, open field test, and stance instability was determined by two-way repeated-measures analysis of variance and Dunnett’s multiple comparison tests against behavioural performance at 0-week time point. Statistical significance for inverted wire hang was determined by two-way repeated-measures analysis of variance and Dunnett’s multiple comparison tests against the unguided condition. We also assayed behaviour in saline-injected controls (not shown) and found them indistinguishable from unguided controls. Points and bars represent mean ± s.e.m. For all groups, *n* = 5 except (ubiquitous + sgCacna1a B) and (crosstalk + sgCacna1a A) groups, in which *n* = 6. \**P* < .05, \*\**P* < .01, \*\*\**P* < .001, \*\*\*\**P* < .0001

As a control for the effects of off-target editing, we used two sequence-independent guide RNAs (sgCacna1a A and sgCacna1a B), comparing these to an unguided condition in which no guide RNA sequence was present. To assess whether this approach could recapitulate known phenotypes resulting from PC-specific loss-of-function of *Cacna1a*, we assessed a battery of behaviours before and for five weeks after AAV administration (Fig. 6a, right side).

In both ubiquitous and PC-specific paradigms, we observed a strong reduction in Cacna1a staining in the cerebellum that was consistent between both guide RNAs (Fig. 6b,c and Extended Data Fig. 8a). Importantly, we saw similar reductions in Cacna1a staining intensity with both ubiquitous and PC-specific SaCas9 expression.

Both ubiquitous and PC-specific *Cacna1a* disruption also recapitulated several hallmarks of ataxia: impairments in skilled motor behaviour, as assessed by narrowing beam crossing (Fig. 6d, Extended Data Fig. 8b and Supplementary Videos 1,2), reduced limb strength (Fig. 6e), reduced locomotion in an open field (Fig. 6f,g and Extended Data Fig. 8c), and gait deficits (Fig. 6h,i and Extended Data Fig. 8d-g). Both guide RNAs resulted in similar phenotypes, suggesting that the deficits observed were not due to off-target editing, and no deficits were observed in animals that did not receive a guide RNA. Whereas ubiquitous expression of SaCas9 led to significantly reduced weight by 3 weeks post-injection, we did not observe any significant difference in weight until 5 weeks post-injection with PC-specific expression (Extended Data Fig. 8h).

These results demonstrate that transcriptional crosstalk can be leveraged for cell type-specific gene manipulation in wildtype animals, with high enough coverage of targeted cells to recapitulate phenotypes from genetic knockouts.

## Discussion

Diversification of the AAV capsid through directed evolution has yielded a toolkit of AAVs with varied tissue tropism^11–23^. Further refinement of expression can be achieved through inclusion of regulatory elements, including enhancer sequences^24–33^. However, successful incorporation of these elements requires an understanding of how AAV genomes are processed by the host cell, necessitating development of novel methods for visualizing and measuring AAV genomes *in situ*.

To this end, we developed two complementary methods: AAV-Zombie and SpECTr (for “<u>Sp</u>ECTr <u>E</u>nables AAV <u>C</u>oncatemer <u>Tr</u>acking”). AAV-Zombie enables single-molecule visualization of the AAV genome *in situ,* allowing us to profile capsid-genome interactions in cultured cells and to assess DNA-level transduction of engineered viral vectors in tissue (Fig. 2). Though protein- and RNA-level measurements of AAV transduction have provided invaluable insights at the single cell level, these methods can miss cell types that are recalcitrant to AAV genome expression^40^. Further refinement of AAV-Zombie, using multiple and/or combinatorial barcodes with a sequential FISH paradigm, may enable DNA-level readout of large capsid pools or libraries *in situ*.

Complementary to AAV-Zombie, SpECTr enables single molecule visualization of AAV concatemers *in situ* (Fig. 3). Development and implementation of AAV gene therapies necessitates better understanding of genome processing in cell types relevant to disease, especially given the stability of episomal AAV genomes in host cells and implicated roles of AAV concatemers in productive transduction^53,64^. SpECTr can help address this critical knowledge gap, as we highlight here with our multiparametric single-cell AAV transduction profiling in primary cell cultures.

Concatemerization of AAV genomes can have unintended consequences when delivering multiple genomes. Duan and colleagues demonstrated transcriptional crosstalk with a ubiquitous enhancer in cultured cells and in muscle tissue^39^. Here, we extend this observation to multiple cell type-specific enhancers and a variety of tissue and cell types. We observed transcriptional crosstalk with all 11 enhancers we tested, suggesting that this is a general phenomenon (Fig. 1). Using SpECTr, we mechanistically link this transcriptional crosstalk to concatemer formation (Fig. 4).

Transcriptional crosstalk between AAV genomes may lead to undesired expression, especially in cases where multiple enhancers are used for simultaneous targeting of different cell types. For example, pooled AAV screening is used to reduce the number of animals necessary to profile enhancer elements^25,31^. Troublingly, crosstalk between genomes in such pools may confound the resulting transduction profiles. Performing pooled AAV enhancer screens in animals with mutations in DNA repair pathways associated with concatemerization may be one solution to this issue. Our results with SCID mice support this approach (Fig. 4).

Transcriptional crosstalk also presents an opportunity for AAV-based expression of large cargo in a cell type-specific manner by separating bulky gene regulatory elements from promoters and coding sequences. We demonstrate this here using Cas9 nuclease, directing editing to specific cell types *in vivo*, with sufficient efficiency and coverage of the target population to recapitulate known loss-of-function behavioural phenotypes (Fig. 6). Importantly, this paradigm does not require transgenic organisms, and so may be easily applied to a variety of models, including disease models where crossing of transgenics would not be feasible, as well as models where Cas9-expressing transgenics are not readily available. This approach is particularly exciting given the recent development of AAV capsids that provide genetic access to the nervous systems of non-human primates following systemic administration^15–17,23,65^ and the identification of gene regulatory elements for cell type-specific expression across species^24,27–29,33^. The small size of minimal promoters and terminator sequences allows easy integration of even large CRISPR effectors, including fusion proteins for AAV-based gene activation^66^, base editing^67^, or prime editing^68^. Targeting of these genome editing tools to specific cell types following minimally-invasive delivery of crosstalking genomes may open new avenues of research into gene function in a myriad of model organisms.

In summary, we identified and profiled transcriptional crosstalk occurring between promoters and cell type-specific enhancers delivered on separate AAV genomes. We paired transcriptional crosstalk with systemically-administered BBB-penetrant AAVs to enable minimally-invasive delivery of a large Cas9 cargo for cell type-specific gene disruption in wildtype animals, which recapitulated phenotypes of genetic knockouts. To understand the mechanisms underlying transcriptional crosstalk, we developed and validated spatial genomics techniques, AAV-Zombie and SpECTr, that enable tracking of AAV genomes and concatemers in intact cells and tissue. Leveraging these methods, we demonstrated that concatemerization of the AAV genome facilitates transcriptional crosstalk. These novel spatial genomics techniques can help to bridge a critical knowledge gap linking AAV genome processing with expression and enable integration of gene regulatory elements for genetic access to and manipulation of targeted cell populations, both in basic research and in translational contexts.

## Methods

### Plasmid DNA

Standard molecular cloning techniques were used to generate DNA constructs in this study. Double-stranded DNA was synthesized by Integrated DNA Technologies and inserted into pAAV backbones with NEBuilder HIFI (New England Biolabs, E2621). sgRNA sequences were synthesized as overlapping single-stranded DNA oligos (Integrated DNA Technologies) that were then annealed together and ligated into sgRNA expression cassettes using T4 DNA ligase (New England Biolabs, M0202).

pUCmini-iCAP-AAV-PHP.eB^12^ (Addgene #175004), pUCmini-iCAP-AAV.CAP-B10^14^ (Addgene #103005), pUCmini-iCAP-AAV.MaCPNS2^15^ (Addgene #185137), AAV-DJ rep-cap (Cell Biolabs, VPK-420-DJ), AAV6 rep-cap (Cell Biolabs, VPK-420-DJ), and pHelper (Agilent, #240071) plasmids were used for production of AAVs. Prior to use, all plasmids were sequence verified via whole-plasmid sequencing (Primordium Labs).

### AAV production

AAVs were produced and purified according to published methods^69^, with some minor alterations. Briefly, HEK293T cells were triple transfected with PEI-MAX (Polysciences, #24765) to deliver the rep-cap or iCAP, pHelper, and genome packaging plasmids. Viruses were harvested from cells and media, then purified over 15%, 25%, 40%, and 60% iodixanol (OptiPrep, Serumwerk, #1893) step gradients. A Type 70 Ti fixed-angle titanium rotor (Beckman Coulter, #337922) at 58.4k rpm for 1.5 hr, or a Type 70.1 Ti fixed-angle titanium rotor (Beckman Coulter, #342184) at 61.7k rpm for 1.25 hr was used, depending on the scale and number of AAVs to be purified simultaneously. Viruses were concentrated using Amicon Ultra-15 or Amicon Ultra-4 filters with a 100 kD size cutoff (MilliporeSigma, UFC9100 and UFC8100) and formulated in sterile DPBS (ThermoFisher, #14190144) with 0.001% Pluronic F-68. AAVs were titered by measuring the number of DNase I-resistant viral genomes, relative to a linearized genome plasmid standard. Prior to injection, AAVs were diluted in sterile saline.

### Tissue culture

For AAV production, and for some *in vitro* experiments, HEK293T cells were used (ATCC, CRL-1573). Cells were grown in DMEM (ThermoFisher, #10569010) supplemented with 10% defined FBS (Cytiva, SH30070.03).

For small-scale HEK293T experiments, cells were seeded at optimal confluence (50% for transduction, 90% for transfection) in the morning, and transfected or transduced in the afternoon. For transfection, Lipofectamine LTX (ThermoFisher, #15338100) was used, with 500 ng total of DNA and 3 μL of transfection reagent. To avoid saturating SpECTr or fluorescent protein signal, 50 ng of each experimental DNA was used, with pUC19 (New England BioLabs, N3041S) used as filler to ensure efficient transfection. For investigation of transcriptional crosstalk with transfection and transduction *in vitro*, we transduced cells with a 1e5 multiplicity of infection (MOI) of AAV-DJ and cells were collected 5 days later. For *in situ* restriction enzyme digest of AAV concatemers, an MOI of 1e6 AAV-DJ was used and cells were collected 3 days later. On the morning of collection, we passaged cells 1:10 onto poly-D-lysine coated coverslips (Neuvitro, GG-12-1.5h-PDL). Once HEK293T cells had attached, the coverslips were washed three times in DPBS and then fixed. For analysis of fluorescent protein expression, cells were fixed with ice-cold 4% paraformaldehyde (PFA, Electron Microscopy Sciences, #15714-S) in 1x PBS for 15 min at 4 °C and stored in 1x PBS at 4 °C until use. For AAV-Zombie or SpECTr, cells were fixed with 3:1 methanol:acetic acid (MAA, Sigma-Aldrich, #322415 and A6283) and stored at -20 °C in 70% ethanol until use.

For primary neuron cultures, coverslips (Neuvitro, GG-12-1.5h-pre) were prepared by coating with poly-D-lysine (0.1 mg/mL overnight, Sigma-Aldrich, P6407), poly-L-ornithine (0.01% overnight, Sigma-Aldrich, P4957), and laminin (0.02 mg/mL overnight, ThermoFisher, #20317015). Primary neurons were prepared by pooling cortices and hippocampi from several E16.5 embryos, and digesting the tissue in 15 U/mL papain (Sigma-Aldrich, P3125). The cell suspension was then treated with DNase I and cells triturated in Hanks balanced salt solution (ThermoFisher, #14025092), with 5% horse serum (ThermoFisher, #16050130), then centrifuged through 4% bovine serum albumin. The cell pellet was resuspended in NeuroCult Neuronal Plating Medium (STEMCELL Technologies, #05713), supplemented with 1:50 NeuroCult SM1 (STEMCELL Technologies, #05711), 0.5 mM GlutaMAX (ThermoFisher, #35050061), and 3.7 μg/mL L-Glutamic acid (Sigma-Aldrich, #49449), and plated at a density of 60,000 per coverslip. At 5 days *in vitro* (DIV), half the media was exchanged for BrainPhys Neuronal Media (STEMCELL Technologies, #05790), also supplemented with 1:50 NeuroCult SM1. For transduction, AAV was diluted in the added growth media. The removed plating media was saved and combined 1:1 with complete BrainPhys media. To minimize prolonged transduction due to AAVs in culture media, we used the 1:1 mix of conditioned plating media and BrainPhys media to perform a complete media change at 3 hr post-transduction, with 3 washes in pre-warmed BrainPhys between the aspiration of the virus-containing media and addition of fresh conditioned media. Subsequently, the media was half-changed with supplemented BrainPhys media every 3 days. Primary neurons were harvested and fixed as described for HEK293T cells above.

### Animals

Animal husbandry and all procedures involving animals were performed in accordance with the Guide for the Care and Use of Laboratory Animals of the National Institutes of Health and approved by the Institutional Animal Care and Use Committee (IACUC) and by the Office of Laboratory Animal Resources at the California Institute of Technology.

8-week old, male C57BL/6J (strain #: 000664), C57BL/6J-background *Prkdc^scid/scid^* (strain #: 001913) and C57BL/6J-background *Rosa26^CAG-LSL-tdTomato^* (strain #: 007914) mice were obtained from the Jackson Laboratory. Mice were housed 3-4 per cage, on a 12 hr light/dark cycle, and had *ad libitum* access to food and water. For behavioural experiments, animals were kept in a reverse light cycle; all behavioural assays were conducted during the dark cycle, between ZT13 and ZT17.

For primary neuron cultures, timed pregnant C57BL/6N dams were obtained from Charles River Laboratories.

### Retro-orbital injection

AAVs were administered via retro-orbital injection during isoflurane anesthesia (1-3% in 95% O_2_/5% CO_2_, provided by nose cone at 1 L/min), followed by administration of 1-2 drops of 0.5% proparacaine to the corneal surface^69^.

### Tissue harvest and processing

Tissue was collected 4 weeks post-AAV administration, except for animals used in the *Cacna1a* knockout behaviour experiment in which tissue was collected 6 weeks post-AAV administration. Animals were euthanized via i.p. injection of 100 mg/kg euthasol.

For AAV-Zombie or SpECTr in tissue, animals were transcardially perfused with 30 mL of ice-cold heparinized 1x PBS, and liver and brain were dissected out. For analysis of fluorescent protein expression, one hemisphere of brain and one lobe of liver were submerged in ice-cold 4% PFA formulated in 1x PBS and fixed overnight at 4 °C. The other hemisphere and another lobe of liver were manually dissected into 1 mm^3^ pieces with regions of interest and flash frozen in O.C.T. Compound (Scigen, #4586) using a dry ice-ethanol bath. O.C.T. blocks were kept at -70 °C until sectioning.

For measurement of viral genomes from bulk DNA, tissue was processed as above, except that unfixed tissue was used immediately for genomic DNA extraction (DNeasy Blood and Tissue Kit, Qiagen, #69504), rather than frozen.

If animals were not used for AAV-Zombie, SpECTr, or bulk DNA extraction, then following perfusion with PBS, animals were perfused with 30 mL of ice-cold 4% PFA in 1x PBS. Relevant tissues were then extracted and post-fixed overnight in 4% PFA in 1x PBS at 4 °C. For sectioning, brain and liver were cryoprotected through immersion in 30% sucrose in 1x PBS. Once the tissue had sunk, it was flash-frozen in O.C.T. Compound using a dry ice-ethanol bath and kept at -70 °C until sectioning.

Sections were obtained using a cryostat (Leica Biosystems). Fixed tissue was sectioned at 80 μm, collected in 1x PBS, and stored at 4 °C until use. Tissue for AAV-Zombie or SpECTr was sectioned at 20 μm, collected on a clean glass slide (Brain Research Laboratories, #2575-plus), allowed to dry, then stored at -70 °C until use.

Immediately prior to imaging, gut tissue and DRGs were optically cleared by overnight room-temperature incubation in RIMS^70,71^, then mounted in RIMS with an iSpacer (SunJin Lab). Gut tissue was cut longitudinally before incubation in RIMS and mounted with the myenteric plexus up.

### Digital Droplet PCR

To measure viral genomes from bulk cortex and liver DNA, digital droplet PCR was used. 1 μg of total DNA was first digested overnight with 20 U of SmaI (New England Biolabs, R0141) at 25 °C, or with 20 U each of KpnI-HF and SpeI-HF (New England Biolabs, R3142 and R3133) at 37 °C. The digests were diluted 1:4, and 5 μL of each dilution was loaded into a 25 μL PCR reaction (Bio-Rad, #1863024). 23 μL of the PCR reaction was used to generate droplets (Bio-Rad, #1863005) on a QX200 Droplet Generator (Bio-Rad). 40 μL of droplets were transferred to a PCR plate, which was sealed with a pierceable heat seal (Bio-Rad, #1814040 and #1814000) and the PCR was run according to the manufacturer’s protocol. Post-PCR, droplets were measured with a QX200 Droplet Reader and analyzed using the QX Manager software (Bio-Rad, #12010213). Double-quenched FAM- and HEX-labeled probe assays (Integrated DNA Technologies) were used to detect GFP sequence and W3SL sequence in the same droplets, and the mean of the two resultant concentrations was used. SmaI and KpnI-HF/SpeI-HF digests yielded similar results; only SmaI digests are shown.

### Immunohistochemistry

IHC, except against Cacna1a, was performed on free-floating sections. For IHC detection of Cacna1a, sections were first mounted onto slides and subjected to heat-induced epitope retrieval by boiling in 1x citrate buffer, pH 6 (Sigma-Aldrich, C9999) for 10 min in a microwave, followed by thorough washing with 1x PBS.

For IHC, sections were blocked in BlockAid Blocking Solution (ThermoFisher, B10710) with 0.1% Triton X-100 (Sigma-Aldrich, #93443). Primary and secondary antibodies were diluted in this blocking buffer. Tissue was incubated with primary antibody overnight at 4 °C and with secondary antibody for 2 hr at room temperature. Following each antibody incubation step, sections were washed 3 times for 10 min each in 1x PBS with 0.1% Triton X-100. For Hoechst labeling, sections were incubated for 10 min with 1/10000 Hoechst 33342 (ThermoFisher, H3570) in 1x PBS, followed by 3 washes in 1x PBS. For segmentation of Purkinje cells, sections were Nissl stained with 1/50 NeuroTrace 435/455 (ThermoFisher, N21479) in 1x PBS, followed by two 1-hr room temperature washes and one overnight wash at 4 °C in 1x PBS with 0.1% Triton X-100. Sections were allowed to dry on slides, and then a coverslip was mounted using Prolong Diamond Antifade Mountant (ThermoFisher, P36965).

The following primary antibodies and dilutions were used: rabbit anti-Cacna1a (1:100, Alomone Labs, ACC-001), chicken anti-GFP (1:1000, Aves Labs, #1020), rabbit anti-TagRFP (for detection of mRuby2, 1:1000, a generous gift from Dr. Dawen Cai, University of Michigan, Cancer Tools, #155266). Fluorophore-conjugated F(ab’)_2_ fragment secondary antibodies (Jackson ImmunoResearch) were used at a 1:1000 working concentration.

### AAV-Zombie and SpECTr of cultured cells

AAV-Zombie and SpECTr protocols, and sequences of Zombie barcodes and their split initiator probes were adapted from Askary *et al.* (2020)^42^. Split initiator probes against endogenous genes and reporter transcripts were designed according to Jang *et al*. (2023)^18^.

For cultured cells on coverslips, a humidified reaction chamber consisting of a 1 mL pipette tip box filled with pre-warmed RNase-free water was used. Parafilm placed on the wafer of the box served as a surface for the *in situ* transcription reaction. Coverslips, previously fixed in MAA and stored in 70% ethanol, were first washed twice in 1x PBS. 20 μL of transcription mixture per coverslip was prepared according to the manufacturer’s protocol (ThermoFisher, AM1334 and AM1330). For simultaneous T7 and SP6 reactions, the T7 buffer was used with 1 μL of each RNA polymerase. For single polymerase reactions, 2 μL of the polymerase was used. 20 μL droplets were pipetted onto the surface of the parafilm. The coverslips were dipped in UltraPure water (ThermoFisher, #10977015), quickly dried by touching their edges to a Kimwipe, then placed cell-side down over the droplets. This reaction was incubated at 37 °C for 3 hr.

Once the transcription reaction was finished, the coverslips were placed cell-side up into a clean 24-well plate and fixed for 20 min at 4 °C with ice-cold PFA in 1x PBS. This was followed by two 5 min washes in 1x PBS, followed by two 5 min washes in 5x SSC (ThermoFisher, AM9770). Samples were then incubated for 15-30 min in pre-warmed probe hybridization buffer, consisting of 2x SSC, 10% ethylene carbonate (Sigma-Aldrich, E26258), and 10% dextran sulfate (Sigma-Aldrich, #3730), at 37 °C. Following this incubation, the coverslips were incubated for 12-16 hr at 37 °C in hybridization buffer plus 2 nM of each probe. Probes for Zombie barcodes, reporter transcripts, and endogenous transcripts were pooled.

After probe hybridization, samples were washed twice for 30 min in stringent wash buffer (2x SSC, 30% ethylene carbonate) at 37 °C, then three times for 15 min in 5x SSC with 0.1 % Tween-20 (Sigma-Aldrich, P1379), and then incubated in HCR amplification buffer (2x SSC, 10% ethylene carbonate) for 20-30 min. HCR hairpins (Molecular Technologies) were heated to 95 °C for 90 s, then cooled to room temperature for 30 min in the dark. For HCR on cultured cells, 30 nM hairpin in amplification buffer was used in a 1-hr amplification reaction. The samples were then washed four times in 5x SSC with 0.1% Tween-20 (10 min per wash, at room temperature).

In some cases, the cytoplasm was labeled with a fluorophore-conjugated poly(dT_30_) probe (Integrated DNA Technologies). Coverslips were incubated with 100 nM of poly(dT_30_) probe in 5x SSC with 0.1% Tween-20 for 1 hr, followed by four 10 min, room temperature washes in 5x SSC with 0.1% Tween-20. Finally, Hoechst 33342 was used to label cell nuclei. Samples were mounted with Prolong Diamond Antifade Mountant.

For co-detection of AAV genomes and capsids, a mouse anti-AAV VP1/VP2/VP3 monoclonal antibody conjugated to Alexa Fluor 488 was used (Clone B1, Progen, #61059-488). Following poly(dT) labeling, the samples were immunolabeled as described above, with an overnight 4 °C incubation with a 1:100 dilution of the primary antibody in blocking buffer.

For *in situ* restriction enzyme digest, coverslips were treated with restriction enzymes after MAA fixation and before *in situ* transcription. Restriction enzyme digests were carried out overnight, at 25 °C for SmaI (New England Biolabs, R0141), and at 37 °C for MscI (New England Biolabs, R0534) and BstEII-HF (New England Biolabs, R3162).

### AAV-Zombie and SpECTr of tissue sections

AAV-Zombie and SpECTr were performed on tissue sections as described above for cultured cells, save for a few differences. Incubations in tissue were performed in a staining tray (Simport, M918), and fixation and washes were done in Coplin jars.

Sliced fresh tissue was first removed from -70 °C storage and allowed to warm to room temperature. Slides were then briefly washed with 1x PBS to remove O.C.T. compound, then fixed for 3 hr in MAA at -20 °C. Residual fixative was washed off with 1x PBS while the transcription mix was prepared. A total of 200 μL of transcription mix was used per slide, which was pipetted onto the slide and spread out with a clean glass coverslip. We found that simultaneous T7 and SP6 transcription in tissue yielded relatively few and small spots from the SP6-driven barcode. Thus, we carried out T7 and SP6 transcription reactions on separate slides. Likewise, T7 RNA polymerase was used at a 1:10 dilution, whereas SP6 RNA polymerase was used at 1:5 dilution. As with cultured cells, *in situ* transcription was carried out at 37 °C for 3 hr.

For the HCR-FISH steps on tissue sections, we used 4 nM of each probe in an overnight 37 °C hybridization. The HCR hairpin concentration was also doubled to 60 nM. Short HCR incubations may result in low signal for endogenous transcripts, whereas long incubations can yield large, unresolvable Zombie barcode spots. Thus, we did an overnight incubation with only hairpins for endogenous transcripts, then switched the amplification solution to one containing all hairpins, for 1 hr.

### Imaging

For imaging of fluorescent protein expression in cultured cells and for obtaining whole section images of mouse brain and liver, a Keyence BZ-X710 epifluorescence microscope was used, with a 10x, 0.45 NA air objective.

For all other imaging, a Zeiss LSM 880 was used. Imaging of fluorescent protein expression and IHC-stained tissue was accomplished with a 10x, 0.45 NA air objective. Imaging of AAV-Zombie and SpECTr signal in cultured cells and in tissue was performed with a 63x, 1.4 NA oil immersion objective. Imaging settings were chosen to capture full dynamic range of the signal without saturating pixels. When possible, laser power was adjusted before adjusting detector gain. Imaging settings were first optimized on control samples, before imaging of experimental samples. Fields of view were chosen while imaging non-experimental channels (e.g. Hoechst or Nissl).

### Image analysis for fluorescent protein and immunohistochemistry samples

For all cell and nuclear segmentation, except segmentation of PCs, Cellpose^72^ was used. Images were batch processed using napari^73^ and the serialcellpose plugin. For HEK293T cells, masks were generated from phase-contrast images. For images of cortex, the fluorescent protein signal was used to generate masks.

PC cell bodies were segmented manually, using the Fiji^74^ distribution of ImageJ, from images of Nissl-stained tissue (Extended Data Fig. 1d). The large size and intense Nissl-staining of the PC cell body, relative to neighboring cells, was used to identify PCs.

For analysis of fluorescent protein intensity in HEK293T cells, cortical cells, and PCs, CellProfiler^75^ (version 4.2.5) was used. Classification of cortical cells and PCs as XFP-positive or XFP-negative was also done using CellProfiler, using empirically determined thresholds based on negative control tissue. For these analyses of cortical cells and PCs, 3 planes (850 μm x 850 μm) from at least 4 non-adjacent sagittal sections were quantified (i.e. at least 12 volumes per animal).

Bulk protein quantification of SCID and WT mice was performed using Fiji, from 3 non-adjacent 100 μm sections per tissue per animal. Cortex and cerebellum were manually segmented from sagittal sections; liver sections were analyzed whole.

To quantify CRISPR-Cas9 editing of the Ai14 locus, tdTomato-positive cells were manually counted using Fiji. Three volumes (850 μm x 850 μm x 64 μm) were captured from each of at least 4 non-adjacent sections per animal. PCs and non-PCs were differentiated based on distinct cell morphology and location.

For analysis of Cacna1a expression in cerebellum, Fiji was also used. Four maximum intensity projections of 850 μm x 850 μm x 30 μm volumes were analyzed per animal. In each image, the molecular layer (ML) and granular layer (GL) were manually segmented, and the total average fluorescence intensity was measured in those regions. For each image, the ML intensity was divided by the GL intensity, and then a per-animal average was determined.

### Image analysis for AAV-Zombie and SpECTr

For analysis of AAV-Zombie and SpECTr spots, segmentation was performed as described above. For primary neurons and HEK293T cells, cell body masks were generated from poly(dT)-TAMRA signal and nuclear masks from Hoechst signal. PC nuclei were manually segmented in Fiji, using large nucleus size, euchromatic nuclear staining, and the presence of *Iptr1* transcript to positively identify PCs.

Quantification and measurement of AAV genomes and concatemers in PCs was accomplished using CellProfiler. Genome and concatemer spots were identified within segmented nuclear masks, using empirically determined spot size thresholds and robust background intensity thresholding, chosen due to the sparse foreground signal.

AAV genomes, concatemers, and capsid puncta were identified in primary neurons and HEK293T cells as described above, with some exceptions. For both HEK293T cells and primary neurons, masks were size filtered, using empirically determined thresholds. Primary neuron masks were further filtered for presence of an overlapping nuclear mask, and a cytoplasmic mask was generated by subtracting the nuclear mask from the cell body mask. EGFP transcript intensity was measured in the entire cell body mask; AAV genome, concatemer, and capsid puncta were quantified in both cytoplasm and nucleus. For HEK293T cells, only nuclear AAV genomes and concatemers were measured.

### Animal behaviour

On each day of behavioural training and data collection, animals were acclimated to the testing room for at least 30 minutes before measurements were taken. Animals were trained on beam crossing and gait measurement assays 1-2 weeks before experimental measurements started. Behaviour equipment was disinfected and deodorized between each animal or, in the case of the open-field test, between each cage.

The open-field apparatus consisted of four square arenas (27 cm x 27 cm), with a camera (EverFocus, EQ700) placed 1.83 m above the floor of the arenas. EthoVision XT 17 (Noldus) was used to capture and subsequently analyze animal locomotion. Each trial consisted of a 2 min habituation period, followed by a 10 min test period. To avoid confounds due to odours from non-cagemates, only animals from the same cage were recorded simultaneously. The average velocity over the course of the experimental period was determined.

The inverted wire hang test was used to measure limb strength^62^. Animals were placed onto a wire mesh screen (6 mm x 6 mm mesh), which was then inverted over the top of a 45 cm tall cylinder with clean bedding in the bottom. A blinded experimenter recorded the latency to fall within a max trial period of 120 s. Three trials were recorded and the average of those three trials used.

To measure skilled locomotion using the narrowing beam assay, a clear plexiglass beam consisting of three 25 cm segments (widths 3.5 cm, 2.5 cm, and 1.5 cm) was elevated above the table surface using empty clean cages, according to published protocol^76^. At the narrow end, an empty cage was placed on its side and bedding from the animal’s home cage placed inside. A white light was also placed over the broad end to motivate animals to move across the beam. For each trial, animals were placed at the end of the widest segment, with all 4 limbs touching the beam surface. Each trial was recorded with a video camera placed to the side and perpendicular to the beam’s length, affording a view of both left and right hindlimbs. A trial was considered complete once the animal had traversed the beam, without turning around, and entered the goal cage. Once an animal had completed three trials, the session was completed. For each trial, a blinded experimenter measured the animal’s time to cross the beam (ignoring time spent paused), and assigned a neurological score^77^: (7) traverses the beam successfully, with no more than 4 foot slips and does not grip the side of the beam, (6) traverses the beam successfully, using hindlimbs to aid in more than 50% of strides, (5) traverses the beam successfully, using hindlimbs to aid in less than 50% of strides, (4) traverses the beam successfully, using a hindlimb at least once to push forward, but without bearing load on limb, (3) traverses beam successfully, by dragging hindlimbs without using them to push forward, (2) moves at least 1 body length, but fails to traverse beam in 120 s trial period or falls off, (1) fails to traverse beam or falls off, and does not move more than 1 body length. The average score and traversal time of the three trials was used for data presentation and statistics.

For gait analysis, we used MouseWalker, according to published protocols for hardware design and analysis^78,79^. A clear acrylic platform, 80 cm long, with a 5.3 cm corridor flanked by 12.5 cm high walls was used. LED lights positioned around the platform enable tracking of animal contacts with the platform surface, through frustrated total internal reflection (fTIR) that is captured using a camera (iPhone 12 Pro, Apple) positioned under the platform. Mice were placed on one end of the corridor, and fTIR recorded as the animal moved across the platform. Animals were recorded until they had completed 3 continuously moving traversals of the field of view. Data were analyzed using MouseWalker.

### Statistics and reproducibility

The number of biological replicates for each experiment are included in the corresponding figure legends. No data were excluded from analyses. Statistical analysis was performed with GraphPad Prism 10.0.3 (GraphPad Software) as described in figure legends.

## Acknowledgements

We thank the entire Gradinaru laboratory for careful review and helpful discussions, especially C. Oikonomou for thorough review of the manuscript. We thank Patricia Anguiano for excellent administrative assistance. We thank Amjad Askary at the University of California, Los Angeles, for help in implementing the Zombie method for detection of AAV genomes. This work was primarily supported by grants from the National Institutes of Health (NIH; Brain Armamentarium UF1MH128336 to V.G) and the CZI Neurodegeneration Challenge Network (V.G.). G.M.C. was supported by a PGS-D from the National Science and Engineering Research Council (NSERC) of Canada.

## Contributions

G.M.C, M.B, and V.G. designed the study. G.M.C. and M.B. designed AAV-Zombie and SpECTr methods. G.M.C. performed molecular cloning, viral production, tissue culture, tissue staining, imaging, behavioural assays, and data analysis. M.B. performed molecular cloning, viral production, tissue staining, and imaging. N.A. performed molecular cloning, tissue culture, viral production, and tissue staining. B.H.B. performed mouse injections, perfusions, histology, tissue staining, imaging, and behavioural characterization. A.M.H.M. and R.A.E. collected and analyzed gait data. E.D.M. prepared primary neuron cultures. S.R.K. contributed invaluable observations of transcriptional crosstalk. X.C. injected mice for characterization of engineered AAV tropism. G.M.C. analyzed data, prepared figures, and wrote the manuscript, with significant input from M.B. V.G. supervised all aspects of this study.

---

**Extended Data Figure 1.**
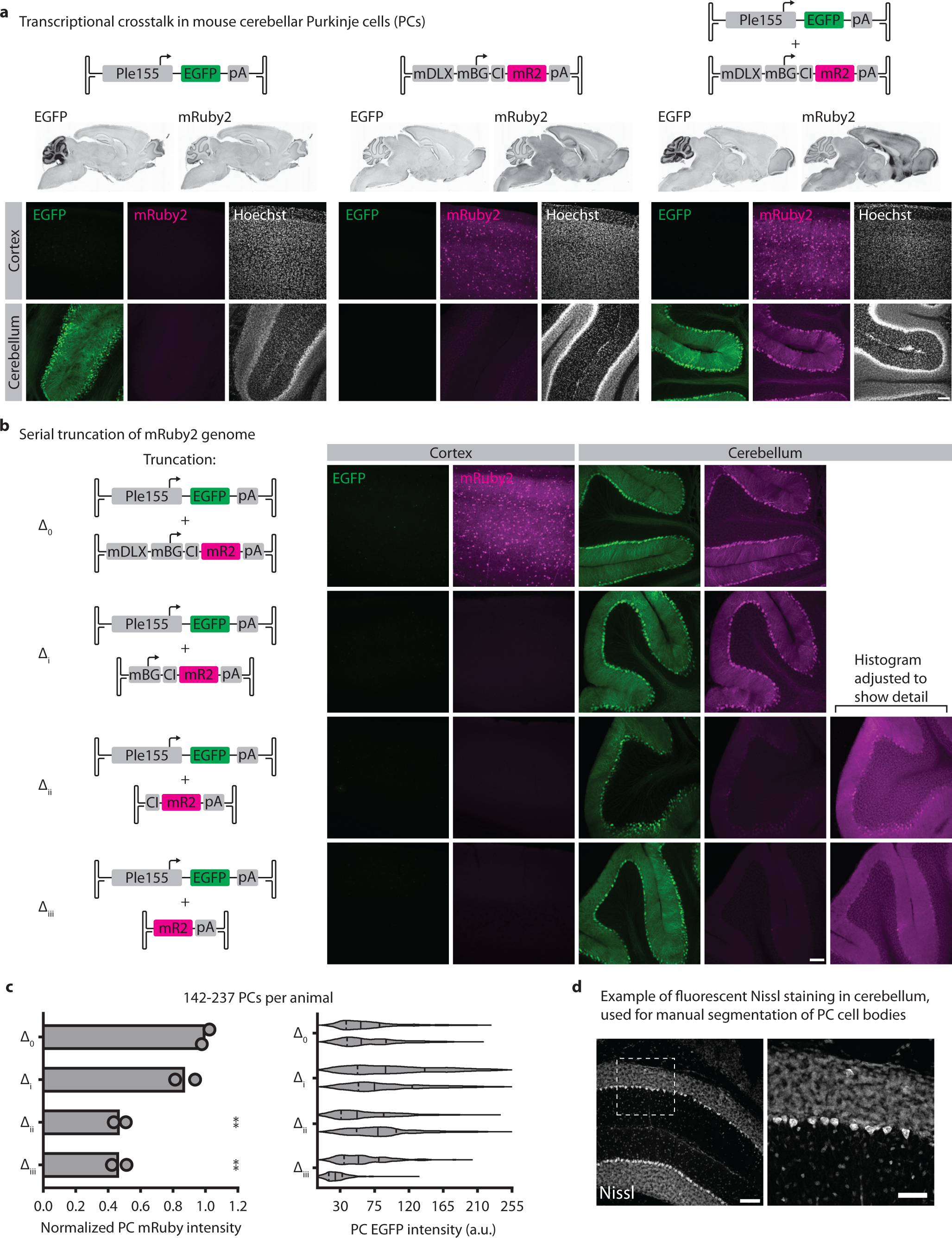
Transcriptional crosstalk in cerebellar PCs between Ple155 and mDLX-minBG. **a**, Related to Fig. 1a,b. Representative images of whole brain, cortical, and cerebellar expression patterns after single or double AAV injections. All genomes delivered at 1e12 vg dose in AAV-PHP.eB. Scale bar = 100 μm. **b,c**, Related to Fig. 1c,d. **b**, Representative images of transcriptional crosstalk between Ple155 element and serially-truncated mDLX-minBG. All genomes delivered at 5e11 vg dose, in AAV-PHP.eB. Scale bar = 100 μm. **c**, Quantification of results for truncation conditions shown in (**b**), quantified as normalized mean PC mRuby2 intensity (left) and PC EGFP fluorescence intensity (right). mRuby2 fluorescence intensity is normalized to the mean of full-length mRuby2 genome (construct Δ_0_). Bars represent mean, with statistical significance determined by one-way analysis of variance and Dunnett’s multiple comparison test against full-length mRuby2 genome. Each violin plot represents EGFP intensity from PCs in one animal. For violin plots, the solid line is the median, and upper and lower dashed lines are quartiles. *n* = 2 animals per condition. All genomes delivered at 5e11 vg dose in AAV-PHP.eB. **d**, Example of fluorescent Nissl staining in cerebellum, showing intense signal in PCs. Nissl staining was used for all manual segmentation of PC cell bodies. Scale bar = 100 μm for left image, 50 μm for right image. CI = chimeric intron.

**Extended Data Figure 2.**
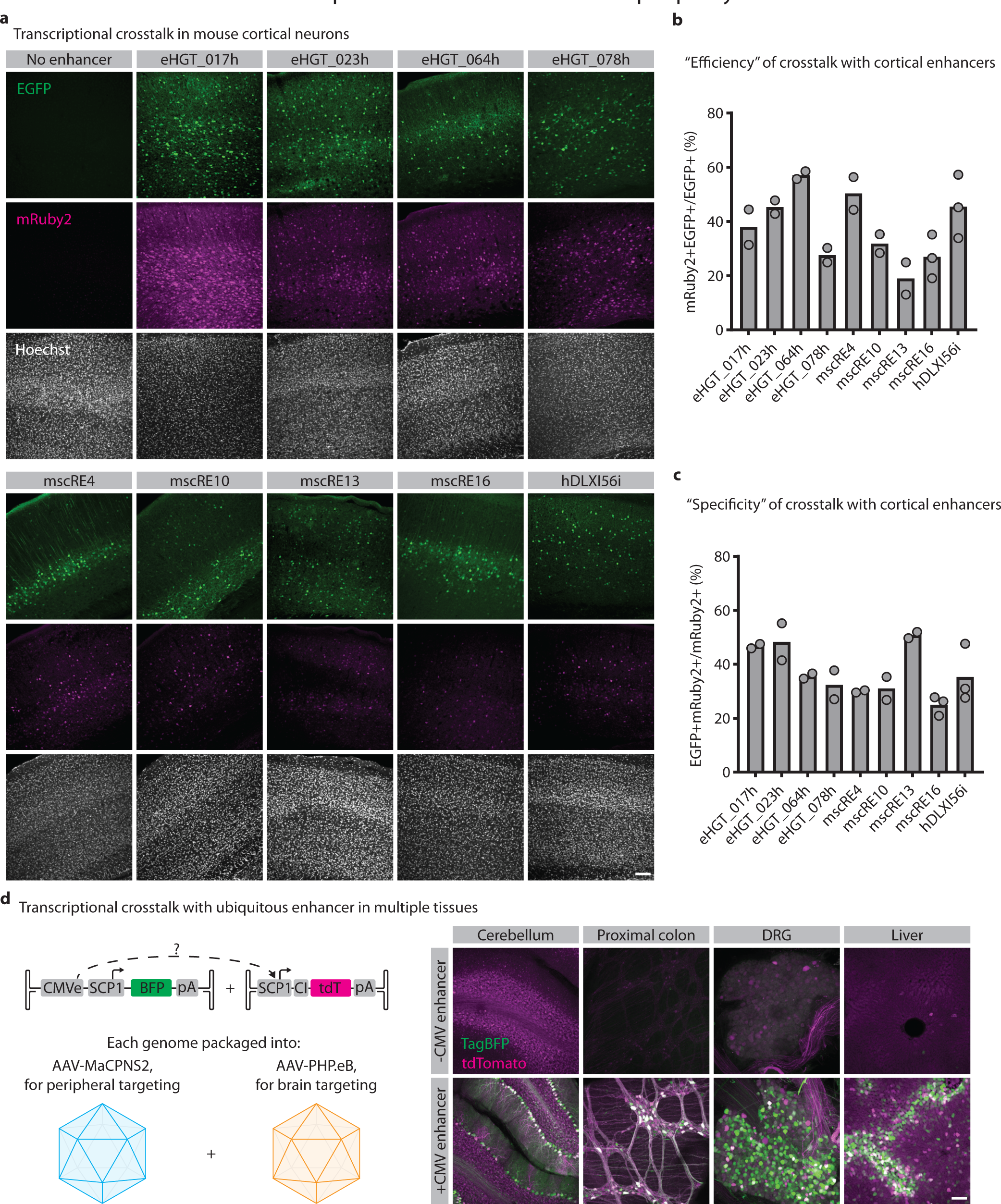
Transcriptional crosstalk in cortex and periphery. **a-c**, Related to Fig. 1e. **a**, Representative images of cortical expression patterns. EGFP reports activity of promoter in *cis* to the enhancer. mRuby2 reports activity of promoter in *trans* to the enhancer. All genomes delivered at 1e12 vg dose in AAV-PHP.eB. Fluorescent protein signal was amplified through IHC. Scale bar = 100 μm. **b**, “Efficiency” of transcriptional crosstalk with various cortical enhancers, quantified as percent of EGFP-positive cells that are also mRuby2-positive. *n =* 2 animals per condition, except mscRE16 and hDLXI56i, for which *n* = 3. **c**, “Specificity” of transcriptional crosstalk with various cortical enhancers, quantified as percent of mRuby2-positive cells that are also EGFP-positive. *n* = 2 animals per condition, except mscRE16 and hDLXI56i, for which *n* = 3. **d**, Transcriptional crosstalk between the ubiquitous CMVe and the SCP1 promoter, in multiple tissues. A cocktail of AAV-PHP.eB and AAV-MaCPNS2 was used to provide broad CNS and PNS coverage (1e12 vg per genome-capsid pair, 4e12 vg per animal). DRG = dorsal root ganglia. Representative images from *n* = 4 animals per condition. Scale bar = 100 μm. Bars in **b** and **c** represent mean.

**Extended Data Figure 3.**
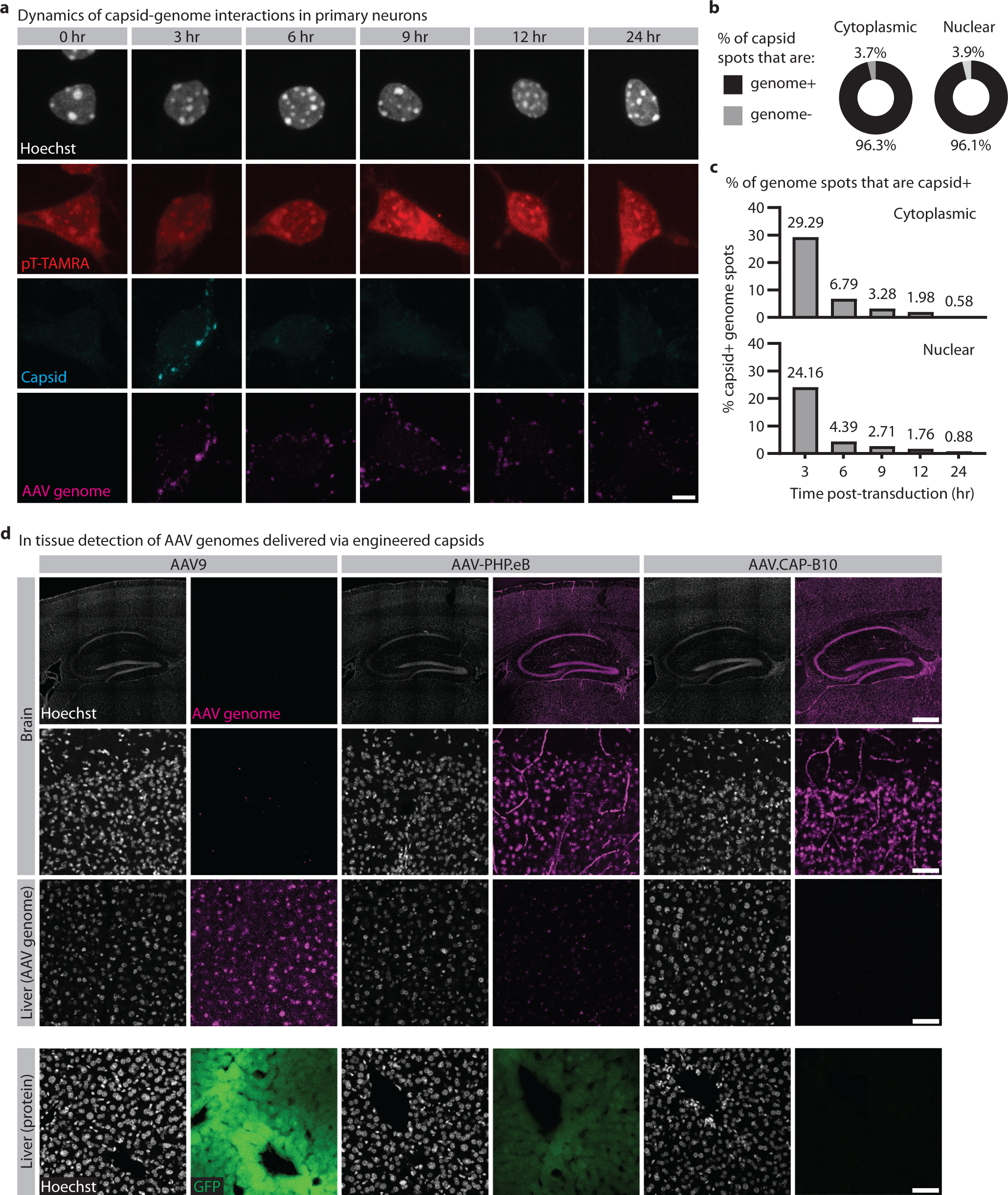
Application of AAV-Zombie to understand AAV transduction *in vitro* and *in vivo*. **a-c**, Related to Fig. 2c. **a**, Representative images of AAV capsids and scAAV genomes, over 24 hrs post-transduction. *n* = 243 (t = 0 hr), 191 (3 hr), 317 (6 hr), 212 (9 hr), 220 (12 hr), 255 (24 hr) neurons per time point. Scale bar = 5 μm. **b**, Percent of capsid puncta that overlap with a scAAV genome, quantified from capsid puncta identified at all time points. The high overlap presumably reflects high encapsidation of the genome. *n* = 763 (nuclear), 1133 (cytoplasmic) capsid puncta. **c**, Percent of genome puncta that overlap with a capsid punctum, for cytoplasmic and nuclear fractions. The decrease in overlap over time reflects uncoating of the AAV genome and degradation of the capsid. For the cytoplasmic fraction, *n* = 3305 (t = 3 hr), 4022 (6 hr), 4201 (9 hr), 5092 (12 hr), 3593 (24 hr) AAV genomes. For the nuclear fraction, *n* = 2421 (t = 3 hr), 3987 (6 hr), 2512 (9 hr), 3469 (12 hr), 2279 (24 hr) AAV genomes. **d**, Related to Fig. 2d. Top 3 rows: AAV-Zombie detection of genomes delivered by engineered capsids AAV-PHP.eB and AAV.CAP-B10, compared to parent capsid AAV9. Tissue was collected 1 day post-injection. Bottom row: EGFP protein in liver, following 2 weeks of expression. Data shows that reduced liver protein expression with AAV-PHP.eB and AAV.CAP-B10 is due to reduced DNA-level transduction, rather than a transcriptional or post-transcriptional difference. Genomes were delivered at 3e11 vg dose. Representative images from *n* = 3 animals per condition. Scale bars = 500 μm for top row, 50 μm for rest.

**Extended Data Figure 4.**
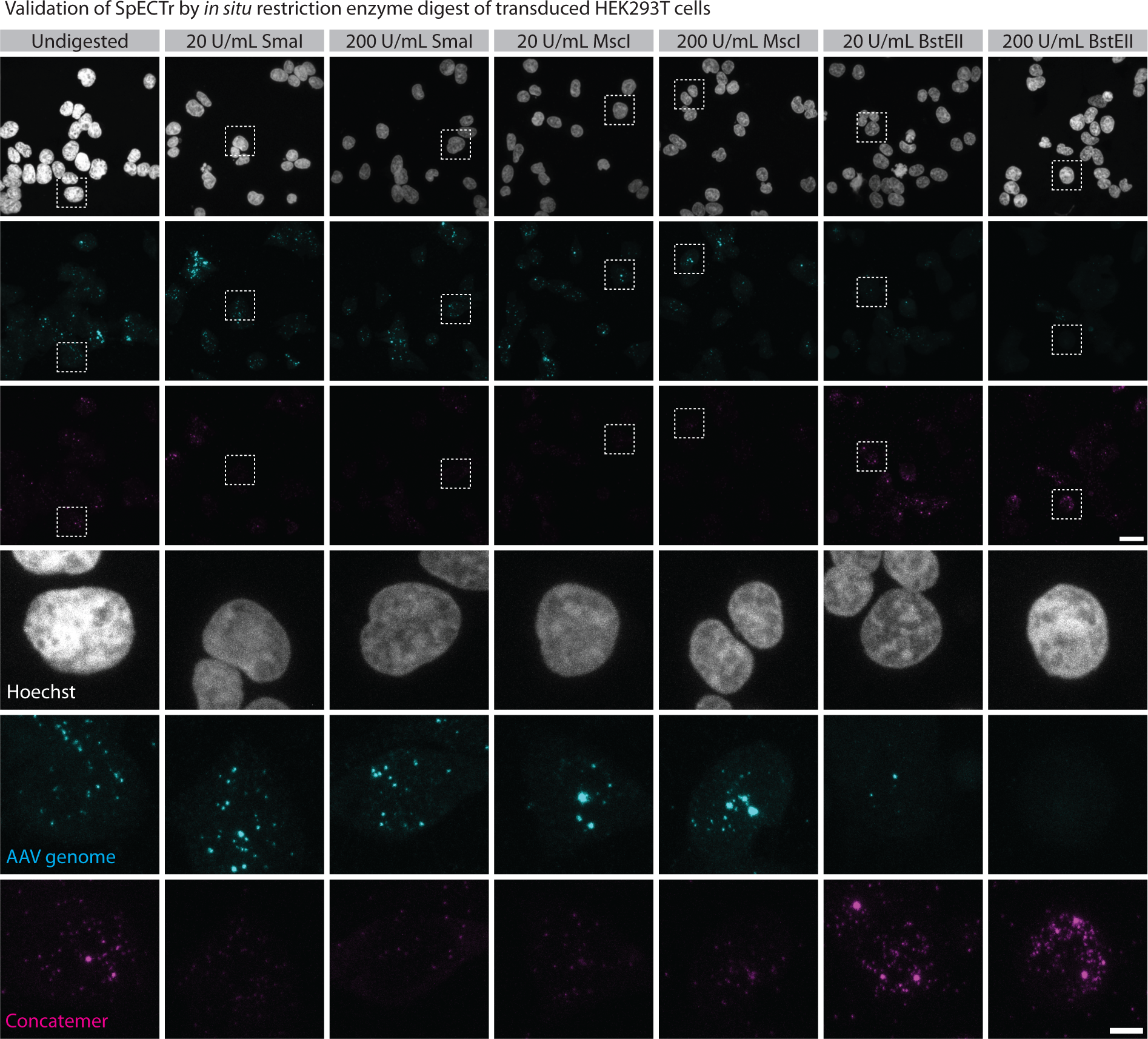
Validation of SpECTr by *in situ* restriction enzyme digest. Related to Fig. 3c-e. Representative images from *in situ* digests of HEK293T cells transduced with SpECTr genomes. “Undig”: undigested condition in which fixed cells were incubated at 37 °C in restriction enzyme buffer without any restriction enzyme. Genomes delivered at 1e6 MOI in AAV-DJ. Scale bars = 20 μm for top 3 rows, 5 μm for rest.

**Extended Data Figure 5.**
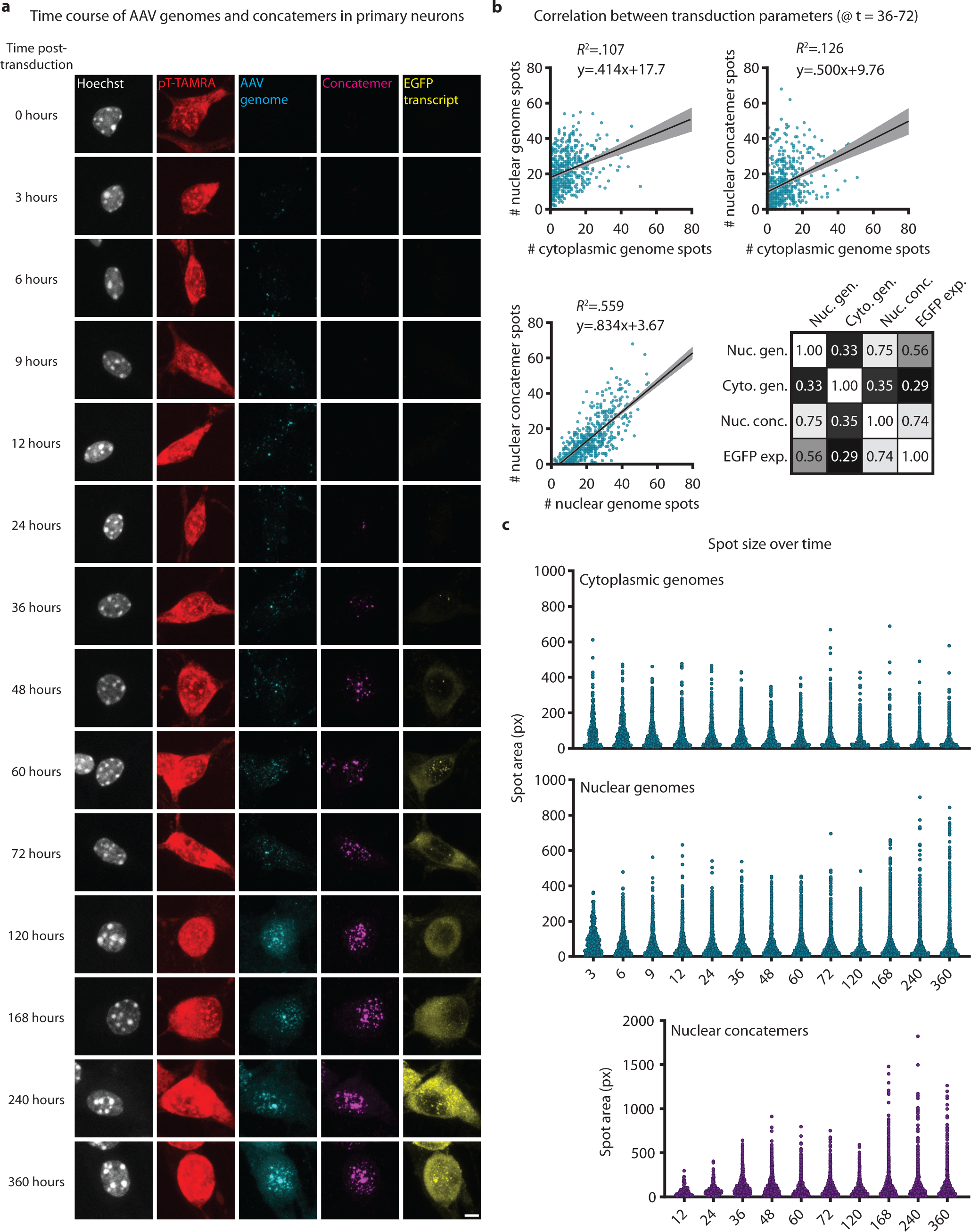
Time course of AAV transduction, concatemerization, and expression in primary neurons. **a-c**, Related to Fig. 3f,g. **a**, Representative images from time course of AAV transduction, concatemer formation, and EGFP reporter transcription in primary neurons. TAMRA-conjugated polyT probe (pT-TAMRA) was used to label cell bodies. **b**, Linear correlations between cytoplasmic AAV genomes, nuclear AAV genomes, and nuclear concatemers, and summary of correlation coefficients for all correlations measured. *n* = 616 primary neurons, pooled from t = 36, 48, 60, and 72 hr time points. **c**, Distribution of spot sizes for cytoplasmic genomes (top), nuclear genomes (middle), and nuclear concatemers (bottom) over time. *n* = 476 - 2098 (cytoplasmic genomes), 657-4078 (nuclear genomes), 111-4226 (nuclear concatemers) spots per time point.

**Extended Data Figure 6.**
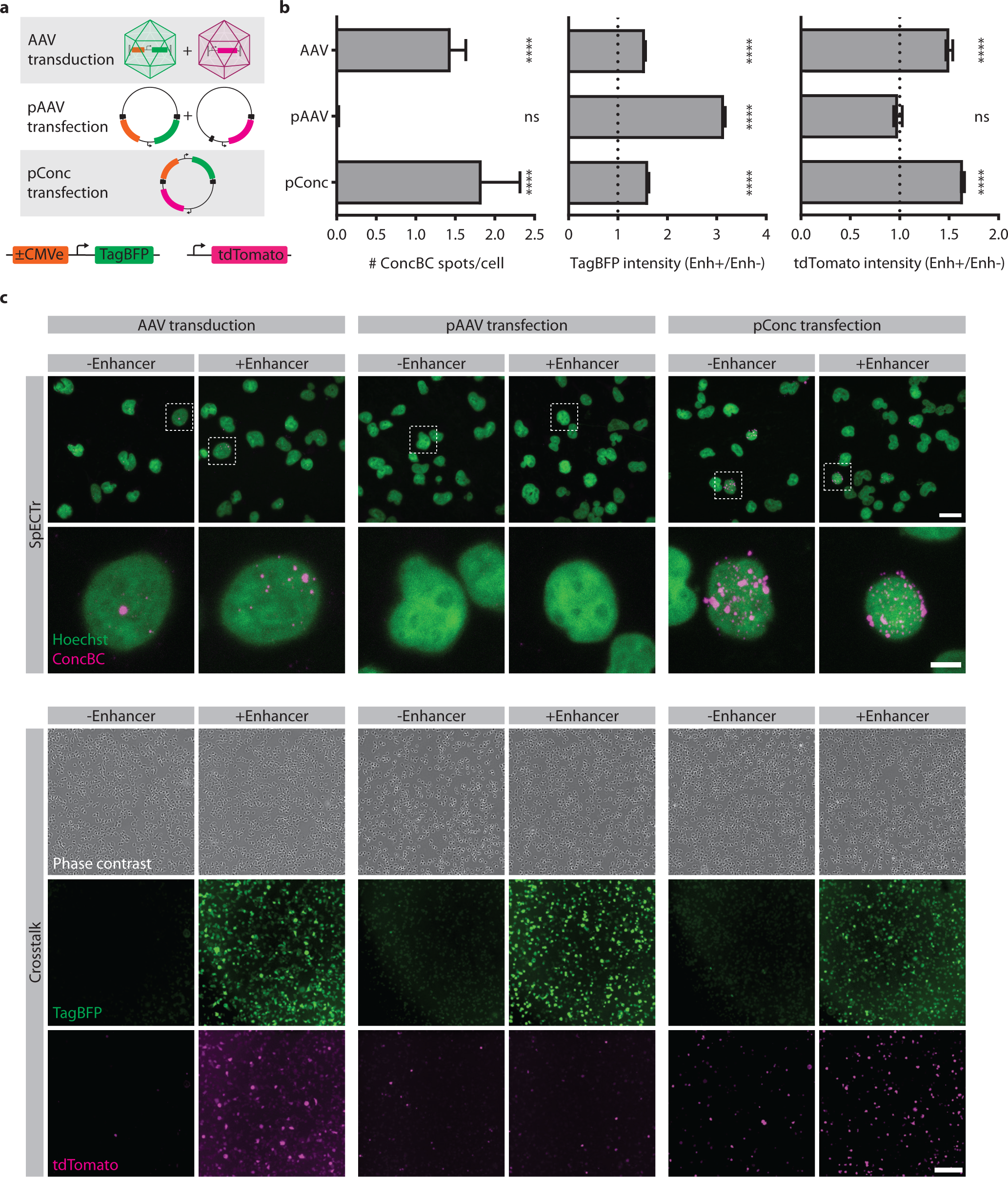
*In vitro* exploration of transcriptional crosstalk mechanisms. **a**, Schematic of experiment. HEK293T cells were either transduced with two cross-talking AAV genomes (top), transfected with two packaging plasmids (middle), or transfected with a single plasmid concatemer (pConc) containing both genomes in *cis* (bottom). Genomes also contained SpECTr elements. **b**, Quantification of SpECTr signal (left), TagBFP intensity (middle), and tdTomato crosstalk reporter intensity (right). Fluorescent protein intensity normalized to the mean of the no-enhancer condition. In all conditions, presence of the enhancer increased expression of TagBFP delivered in *cis*. However, presence of the enhancer increased expression of the tdTomato crosstalk reporter only in the AAV transduction and pConc transfection conditions. Statistical significance for SpECTr signal was determined using a Wilcoxon signed rank test, against the null hypothesis that spot count = 0. *n* = 190 (AAV transduction), 220 (pAAV transfection), 227 (pConc transfection) HEK293T cells per condition. Statistical significance for fluorescent protein intensity was determined using one sample t-tests, against the null hypothesis that normalized intensity = 1. For TagBFP: *n =* 14334 (AAV transduction), 18773 (pAAV transfection), 11606 (pConc transfection) TagBFP-positive HEK293T cells per condition. For tdTomato: *n =* 2675 (AAV transduction), 696 (pAAV transfection), 11613 (pConc transfection) tdTomato-positive HEK293T cells per condition. **c**, Representative images for data quantified in (**b**), showing SpECTr signal (upper panel) and reporter fluorescence (lower panel). Scale bars = 20 μm for top row, 5 μm for second row, and 100 μm for rest. \*\*\*\**P* < .0001, ns = not significant.

**Extended Data Figure 7.**
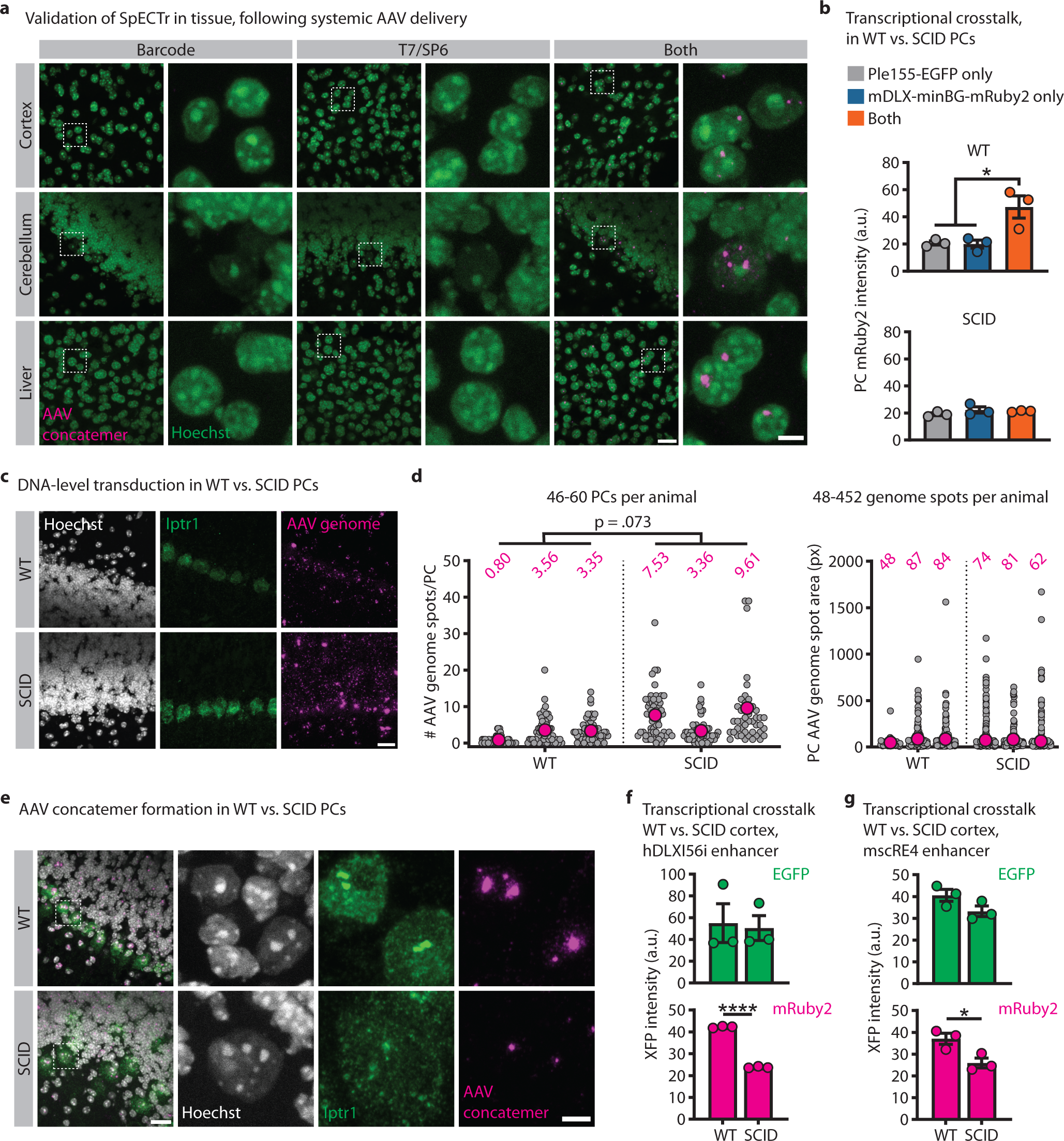
AAV concatemer formation and transcriptional crosstalk in WT and SCID animals. **a**, Validation of SpECTr for detecting AAV concatemers in tissue. Wildtype C57BL/6J mice were transduced with only the barcode genome (left 2 columns), only the T7/SP6 genome (middle), or both genomes (right). AAV concatemers were nuclear and only detected with SpECTr following co-transduction of both genomes. Scale bars = 20 μm for left images of pairs, 5 μm for right. All genomes delivered at 3e11 vg dose in AAV-PHP.eB. **b**, Related to Fig. 4a,b. Quantification of transcriptional crosstalk in WT and SCID PCs, quantified as mean PC mRuby2 fluorescence intensity. Statistical significance was determined using one-way analysis of variation and Tukey’s multiple comparison test. *n* = 3 animals per condition. Bars represent mean ± s.e.m. **c-e**, Related to Fig. 4c,d. **c,** Representative images of AAV genomes detected and quantified with AAV-Zombie in PCs of dual AAV-injected WT and SCID animals shown in Fig. 4a. HCR-FISH against *Iptr1* transcript serves as a marker for PCs. Scale bar = 20 μm. **d**, Quantification of PC AAV genome spot count (left) and spot size (right), in WT and SCID PCs. Each grey dot corresponds to a single PC (left) or a single genome spot (right). Magenta dot and number indicate mean of animal. *n* = 3 animals per condition. Statistical significance was determined using unpaired t-tests. **e**, Representative images of AAV concatemers detected with SpECTr in PCs of dual-injected WT and SCID animals shown in Fig. 4a. HCR-FISH against *Iptr1*transcript serves as a marker for PCs. Scale bars = 20 μm for left column, 5 μm for others. **e,f**, Related to Fig. 4e-h. Quantification of transcriptional crosstalk in WT and SCID cortical cells, quantified as mean XFP fluorescence intensity, with hDLXI56i enhancer (**e**) and mscRE4 enhancer (**f**). Statistical significance was determined using unpaired t-tests. \**P* < .05, \*\*\*\**P* < .0001

**Extended Data Figure 8.**
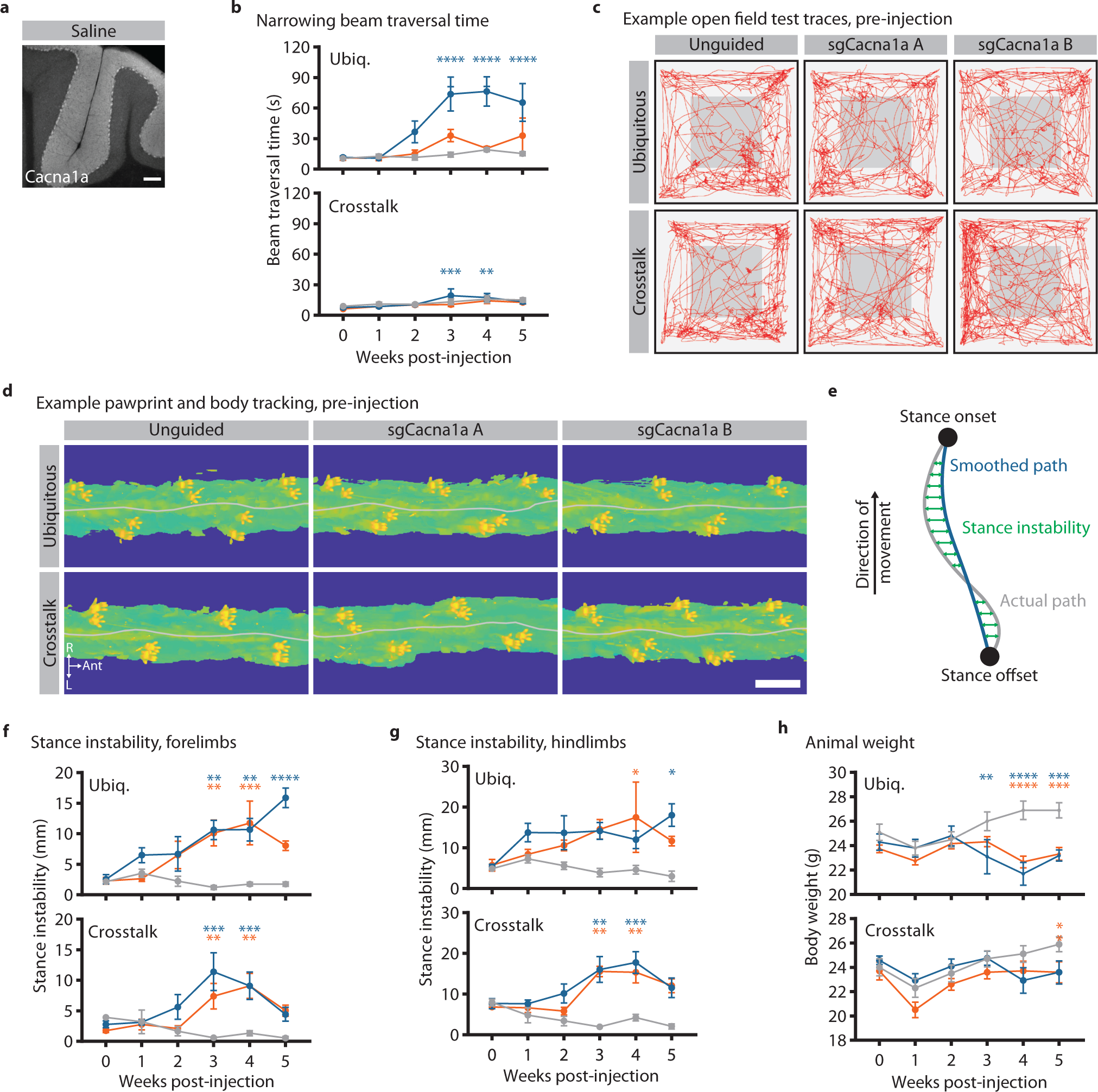
Ataxic phenotypes resulting from transcriptional crosstalk-mediated knockout of Cacna1a in Purkinje cells. **a**, Related to Fig. 6b,c. Representative image of IHC against Cacna1a in cerebellum of saline-injected control animal. Scale bar = 100 μm. **b**, Related to Fig. 6d. Beam traversal time for narrowing beam assay following ubiquitous and PC-specific disruption of *Cacna1a*. **c**, Example open field test traces, acquired pre-injection. Red lines represent animal position over a 10 min trial. **d**, Example pre-injection pawprint and body tracking, over a small segment of the elevated plexiglass platform used for gait analysis. Grey line indicates midline of body. Scale bar = 3 cm. **e**, Related to Fig. 6h,i. Schematic to demonstrate stance instability metric. For each stance, tracking of body center relative to paw location yields a stance trace (grey line) and a smoothed version of that trace (blue line). The difference between the actual path and the smoothed path corresponds to the stance instability. **f**,**g**, Related to Fig. 6h,i. Stance instability for forelimbs (**f**) and hindlimbs (**g**) following ubiquitous and PC-specific disruption of *Cacna1a*. **h**, Animal weights following ubiquitous and PC-specific disruption of *Cacna1a*. Statistical significance for beam crossing and stance instability was determined by two-way repeated-measures analysis of variance and Dunnett’s multiple comparison tests against behavioural performance at 0-week time point. Statistical significance for animal weight was determined by two-way repeated-measures analysis of variance and Dunnett’s multiple comparison tests against the unguided condition. Points and bars represent mean ± s.e.m. For all groups, *n* = 5 except (ubiquitous + sgCacna1a B) and (crosstalk + sgCacna1a A) groups, in which *n* = 6. \**P* < .05, \*\**P* < .01, \*\*\**P* < .001, \*\*\*\**P* < .0001

**Supplemental Figure 1.**
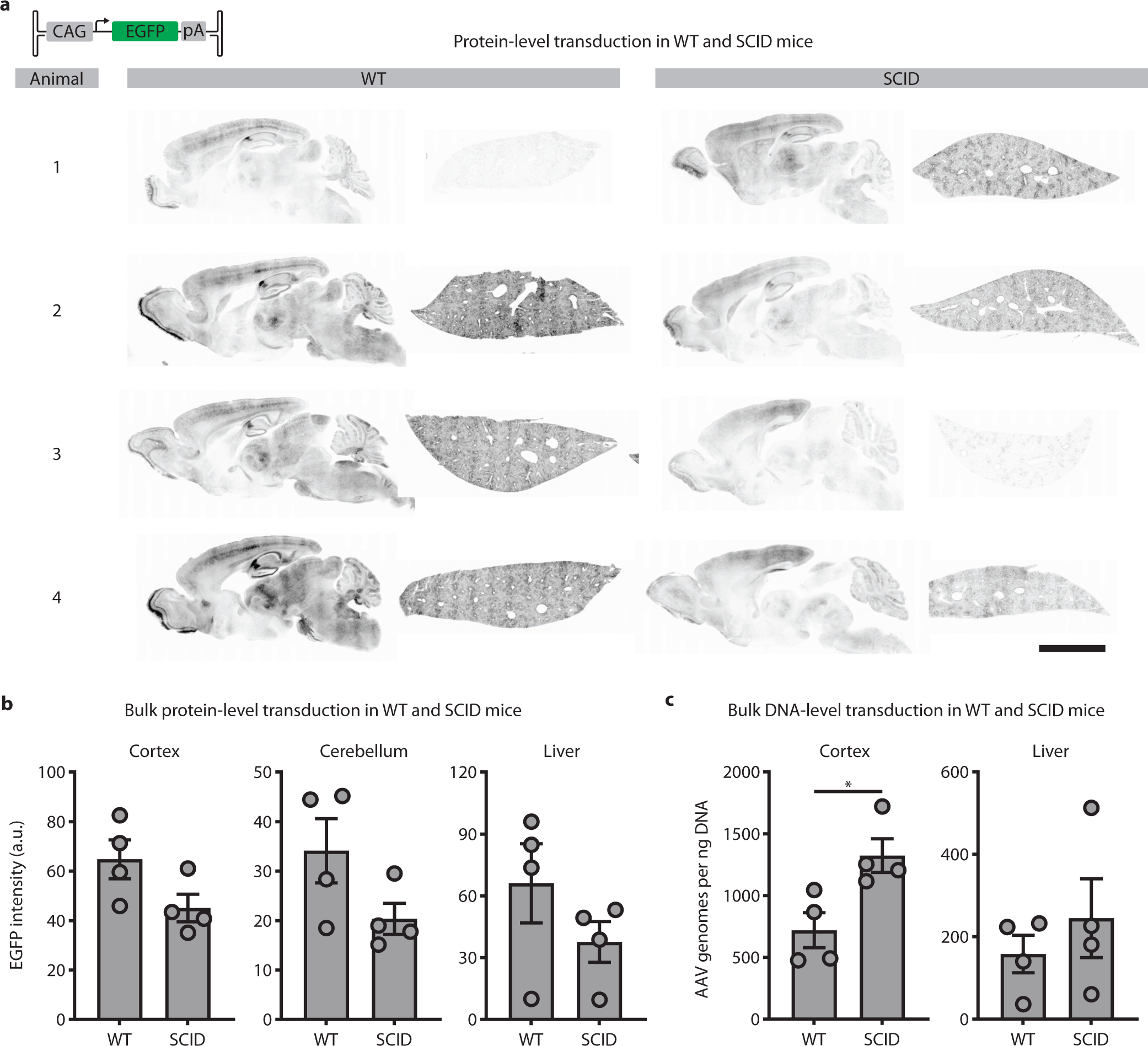
AAV transduction of wildtype and SCID mice. **a**, Wildtype and SCID C57BL/6J animals were transduced with 3e11 vg of AAV-PHP.eB packaging a CAG-EGFP reporter, and tissue was collected 4 weeks later. Representative sagittal brain (left) and liver (right) sections are shown. Scale bar = 5 mm. **b**, Quantification of bulk protein in cortex (left), cerebellum (middle), and liver (right) of WT and SCID animals, assessed by mean EGFP intensity. **c**, Quantification of bulk viral DNA in WT and SCID cortex and liver, assayed through digital droplet PCR. SmaI digests and KpnI-HF/SpeI-HF digests yielded similar results; results from SmaI-digested samples are shown. For (**b**) and (**c**), statistical significance was determined using unpaired t-tests. \**P* < .05

**Supplementary Video 1. Representative videos of narrowing beam crossing performance for animals in ubiquitous SaCas9 condition.** For display purposes, videos are trimmed to show crossing of 2.5 mm segment of beam. The entire length of beam was used for data analysis. Videos show the same animals pre-injection and 4 weeks post-injection.

**Supplementary Video 2. Representative videos of narrowing beam crossing performance for animals in crosstalk-mediated PC-specific SaCas9 condition.** For display purposes, videos are trimmed to show crossing of 2.5 mm segment of beam. The entire length of beam was used for data analysis. Videos show the same animals pre-injection and 4 weeks post-injection.

